# Wound closure after brain injury relies on force generation by microglia in zebrafish

**DOI:** 10.1101/2024.06.04.597300

**Authors:** Francois El-Daher, Louisa K. Drake, Stephen J. Enos, Daniel Wehner, Markus Westphal, Nicola J. Porter, Catherina G. Becker, Thomas Becker

## Abstract

Wound closure after a brain injury is critical for tissue restoration but this process is still not well characterised at the tissue level. We use live observation of wound closure in larval zebrafish after inflicting a stab wound to the brain. We demonstrate that the wound closes in the first 24 hours after injury by global tissue contraction. Microglia accumulation at the point of tissue convergence precedes wound closure and computational modelling of this process indicates that physical traction by microglia could lead to wound closure. Indeed, genetically or pharmacologically depleting microglia leads to defective tissue repair. Live observations indicate centripetal deformation of astrocytic processes contacted by migrating microglia. Severing such contacts leads to retraction of cellular processes, indicating tension. Weakening tension by disrupting the F-actin stabilising gene lcp1 in microglial cells, leads to failure of wound closure. Therefore, we propose a previously unidentified mechanism of brain repair in which microglia has an essential role in contracting spared tissue. Understanding the mechanical role of microglia will support advances in traumatic brain injury therapies

**Graphical Abstract:** 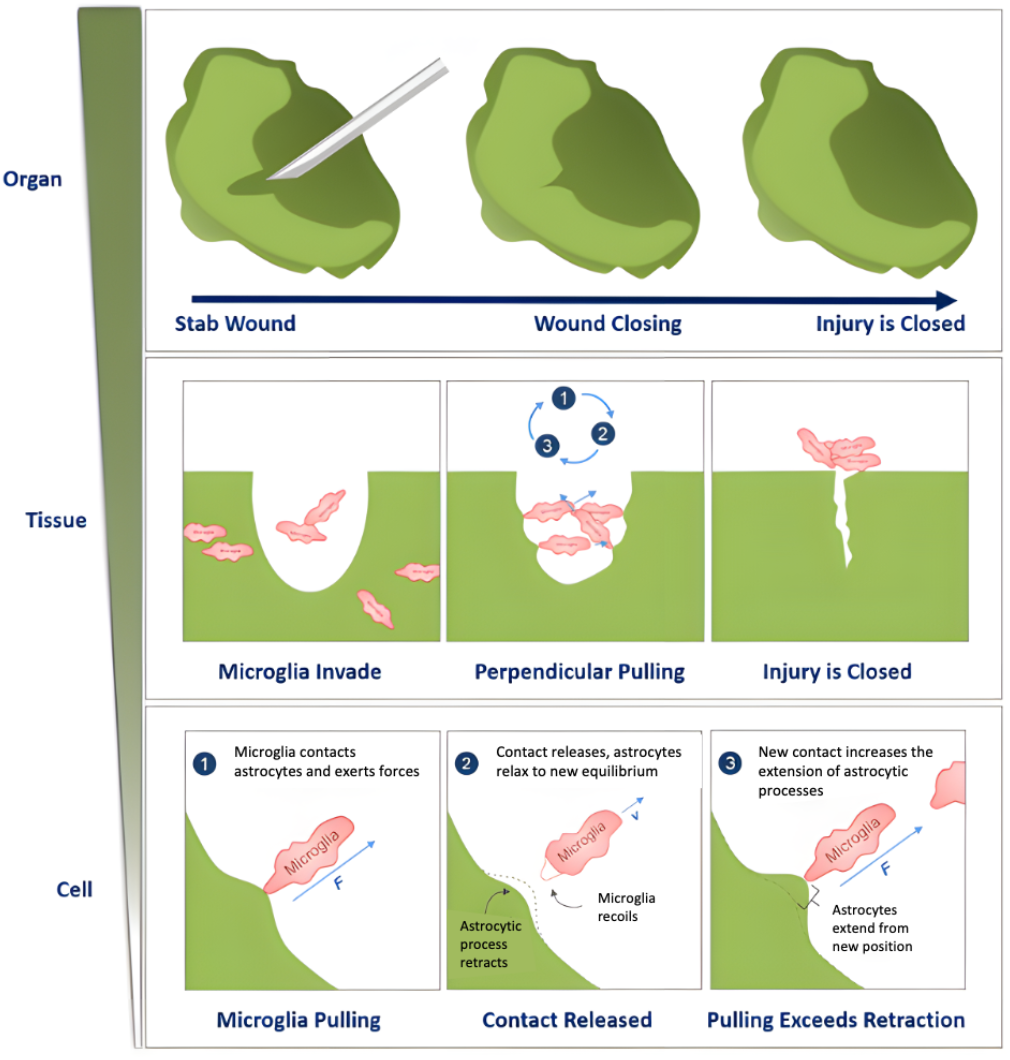

## Introduction

Traumatic brain injury (TBI) is a severe condition affecting an increasing number of people, estimated from 27 million to 69 million each year [1, 2]. Brain trauma often leads to serious health problems including persistent impairments in cognitive, sensory, and motor function. Unfortunately, there is no treatment yet for this complex condition and brain tissue in mammals has a very limited capacity to repair. In contrast, zebrafish larvae possess high regenerative capacities and can fully repair brain injuries within a few days [3]. They are optically transparent which allows for real-time *in vivo* visualisation of tissular and cellular processes. Moreover, these fish share a high degree of similarity in genetics, neuroanatomy and social behaviour with humans. This makes them an attractive animal model for TBI studies [4]. In particular, little is known about the early cellular responses to TBI. One distinct feature of penetrating TBI is the formation of a wound. Understanding wound closure and the factors affecting its dynamics could bring important insights for therapeutic applications. Recovery of brain tissue integrity could be achieved by insertion of new neurons and glial cells or by surviving neurons migrating to the injury site and filling the wound [5, 6]. Microglia, the resident immune cells of the central nervous system (CNS), are the first cell type to respond to a brain injury and home in on the damaged tissue [3, 7, 8]. They are recruited where the tissue is damaged, sequester chemicals, and engulf dead cells and cellular debris. Microglia are also known to interact with many cells, in particular neurons and astrocytes in the injured brain [9]. Interestingly, microglial cells can also exert physical forces [10] and could contribute to would closure by pulling on surrounding tissue. Therefore, these cells are a primary target for investigating their roles in brain injuries [11].

In a model of stab injury in the optic tectum, we show that such wounds close within 24 hours. Tissue closes via large tissue deformations and not through migration or proliferation of neural cells. Live imaging and mathematical modelling suggest that mechanical forces from accumulating microglia can contribute to wound closure by pulling on astrocytic processes. Astrocyte/microglia contacts are under tension as shown by laser ablation. Furthermore, genetically and pharmacologically preventing microglia accumulation as well as destabilising F-actin in microglia, by disrupting lcp1, all inhibited tissue closure. These observations indicate a previously unrevealed mechanical role of microglia in wound healing and inflammatory response after TBI. Overall, we propose a new biophysical perspective on tissue repair and the action of immune cells in the early stages of brain restoration after TBI in which mechanical forces could have a pivotal role.

## Results

### Optic tectum wounds in injured brains close by tissue contraction

Investigating how wounds close after an injury is crucial to our understanding of how the brain can be efficiently repaired. We used an experimental paradigm of stab injury to the optic tectum in zebrafish larvae previously described [3]. In this paradigm the only immune cells in the brain parenchyma are microglia, identified by expression of a transgenic reporter for p2ry12 and expression of the 4C4 antigen [3]. The system thus offers the opportunity to observe the microglia’s reaction to a brain wound in the absence of neutrophils, blood-derived macrophages and cells of the adaptive immune system, which only develops later. To inflict the lesion, we inserted an insect pin at an angle into the middle of the optic tectum, avoiding peripheral proliferation zones and targeting areas where neurons have a laminar and columnar arrangement (Fig. 1A).

**Fig. 1.**
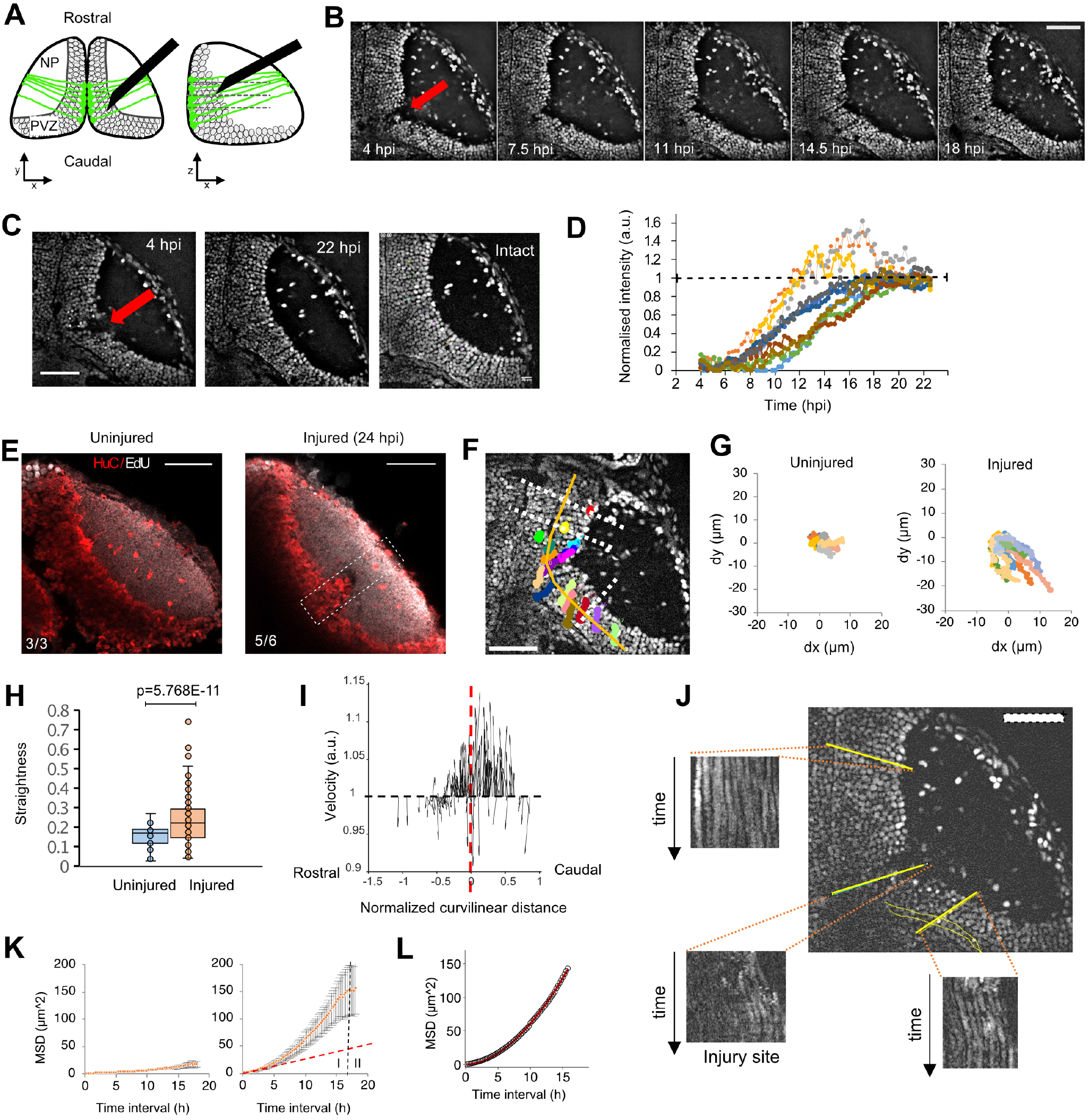
The optic tectum wound closes within 24 h after the injury. (A) Organisation of neurons in the optic tectum and our injury model. Neurons (dark circles), surrounded by astrocytic processes (green) are arranged in layers (lamina, dashed lines) and columns as shown in XY and XZ sections, an insect pin (black) is used to injure the optic tectum down to the periventricular zone. NP: neuropil, PVZ: periventricular zone. (B) Time series of Tg(h2a:GFP) 4 dpf larvae after an injury showing wound closure (red arrow indicates wound position). (C) Wound closure between 4 hpi (left) and 22 hpi (middle) shows the restoration of the tissue as compared to the intact tectum (right; red arrow indicates wound position). (D) Variation over time of the fluorescence intensity inside the injury site during tectum repair (n = 11 larvae). (E) EdU staining (white) and HuC immunofluorescence (red) in Tg(h2a:GFP) uninjured larvae (left) and injured (right) larvae. (F) Individual trajectories of neuron nuclei in the tectum of an injured Tg(h2a:GFP) 4 dpf larvae (coloured traces). The starting points of individual trajectories were projected perpendicularly (white lines) on the curved rostro-caudal extend (orange curve) (G) Individual trajectories in uninjured and injured larvae showed an increased and directional XY displacement for injured animals. (H) Straightness analysis of uninjured (n=7) and injured (n=11) larvae showing that trajectories of neuronal nuclei are more elongated after injury. Box plots show the median, box edges represent the 25th and 75th percentiles, and whiskers indicate ± 1.5 x the interquartile range. (p-value <0.0001, t-test) (I) Trajectory directionality analysis of individual nuclei after rostro-caudal curve linearisation indicates trajectory anisotropy. The injury centre position is represented by the dashed red line. (J) Kymographs at different locations in the PVZ area of an injured fish show that neurons keep their laminar organisation over time. (K) Mean-Square Displacement (MSD) analysis of individual trajectories of neuronal cell nuclei for uninjured (left) and injured (right) larvae show two phases of displacement: I, a super-diffusive behaviour after injury, II, a slower increase compatible with diffusion. The red dashed line shows the MSD for a diffusive case. (L) MSD super-diffusion model, *MSD* (*t*) ∝ *t*^*β*^, fit of the experimental MSD curve in d. blue curve: theoretical model, red points: experimental values. R> 0.999, *β*= 1.86. All images oriented with the rostral side up. Scale bars represent 50 µm on all images.

The optic tectum is located in the upper part of the midbrain which makes it an ideal target for penetrative injuries and live imaging. We took care when performing the direct lesion injury not to harm other parts of the brain, while still forming a clear wound in the periventricular zone (PVZ) initiating a microglial response.

To analyse the kinetics of wound closure, we injured 4 days post-fertilisation (dpf) Tg(h2a:GFP) zebrafish larvae in which cell nuclei were fluorescently labeled. We then performed 3-dimensional time-lapse imaging between 4 and 22 hours post-injury (hpi; Fig. 1B,C; Movie S1). We measured the GFP fluorescence intensity normalised between 0 (before wound closure) and 1 (total intensity in the wound at 22 hpi) over time in the median plane of time series inside the injured region of the periventricular zone. Here the wound is most clearly delimited because of the compact arrangement of neuronal cell bodies (Fig. S1A-D). Our data show that the stab wound is closed (i.e. filled by cells) between 18 and 22 hpi with similar kinetics among different animals (Fig. 1D). Hence, robust wound closure is achieved in less than 24 hpi.

To investigate the mechanisms leading to wound closure, we first investigated whether cell proliferation can lead to the repopulation of the injury site. To detect newly generated neurons and other potential mitotic cells, we applied EdU from the time of injury in combination with immunofluorescence for neurons (HuC) and counted the newborn cells at 24 hpi. We detected labeled cells in the peripheral growth zones of the tectum, which served as an internal control for the EdU staining. However, in the injury site, there were no EdU or EdU/HuC double-labelled cells in 5 out of 6 injured animals. In 1 fish, we observed 4 EdU/HuC double-labeled cells, and 6 EdU-only labeled cells in a superficial position that were probably displaced during the tissue preparation. We also did not observe any EdU+ cells in a corresponding position in 3 uninjured fish (Fig. 1E). In time-lapse recordings of labelled neuronal cells in Tg(Xla.Tubb:DsRed) we also did not observe pre-existing neurons migrating into the injury site (Movie S2) [12]. In some preparations, we noticed an increased number of HuC/D positive cells in the neuropil region where the pin damaged the tissue. However, this likely represents extruded material from the periventricular area, caused by the removal of the pin during lesioning. Additionally, Herzog et al. showed, using the same injury method, that dead cells are present in the neuropil at 6 hpi [3]. Hence, wound closure in our experimental paradigm was not due to neurons filling in the injury site. Instead, we propose that whole tissue movement closes the wound.

To characterise such movement, we followed individual neuronal cell nuclei in discrete locations in the ventricular cell layer by marking their nuclei using Tg(h2a:GFP) larvae in time-lapse recordings (Fig. S2). These neurons are in a columnar arrangement and densely interwoven with astrocytic processes and can therefore be used to track tissue movement. After stab injury, tracked neurons globally showed a large displacement, whereas in intact animals there was minimal movement (Fig. 1F,G). Also, the trajectories of cells in the injured animals were relatively straight, compared to the small and random displacement of neurons in intact fish (Fig. 1H). In injured fish, neurons located in the most caudal part of the tectum experienced a large displacement towards the neuropil. In contrast, those located on the rostral side showed a small lateral displacement toward the opposite hemisphere (Fig. 1I). Kymograph analysis showed that neighbouring cells moved at similar speeds in parallel trajectories without overtaking each other (Fig. 1J), which is expected for uniform tissue movement. Furthermore, we quantified in more detail cellular movements using 2D mean square displacement (MSD) analysis [13] (Fig. 1K). The analysis of injured zebrafish showed two phases, phase I being a super-diffusive behaviour with a power exponent of 1.86 (Fig. 1l; 4 hpi – 20 hpi). Phase II (20 hpi – 22 hpi) indicates that the displacement slows down at longer times (> 20 hpi), correlated with the plateau of the wound closure kinetics curve after 19 hpi (Fig. 1D).

To further characterise putative tissue movements, we determined changes in the outline of the neuropil and ventricular border over time by performing time-lapse experiments where the animals were continuously imaged. We then analysed images from specific time points (typically first and last) from the time series. When required, images were registered for 3D motion as detailed in the Material and Methods section. In this part of the brain the finely branched processes of specific cells, called ependymoradial glial cells or radial glia, are positive for the astrocyte marker glial fibrillary acidic protein (GFAP) and fulfil all astrocytic functions in homoeostasis and after injury [14]. For that reason, we call these processes astrocytic processes.

For visualising the neuropil which comprises both neurites and astrocytic processes, we used Tg(her4.3:GFP-F);Tg(elavl3:MA-mKate2) double transgenic animals, labelling the dense network of astrocytic processes and neuronal membranes with the membrane-tethered fluorophores GFP and mKate2, respectively (Fig. 2A).

**Fig. 2.**
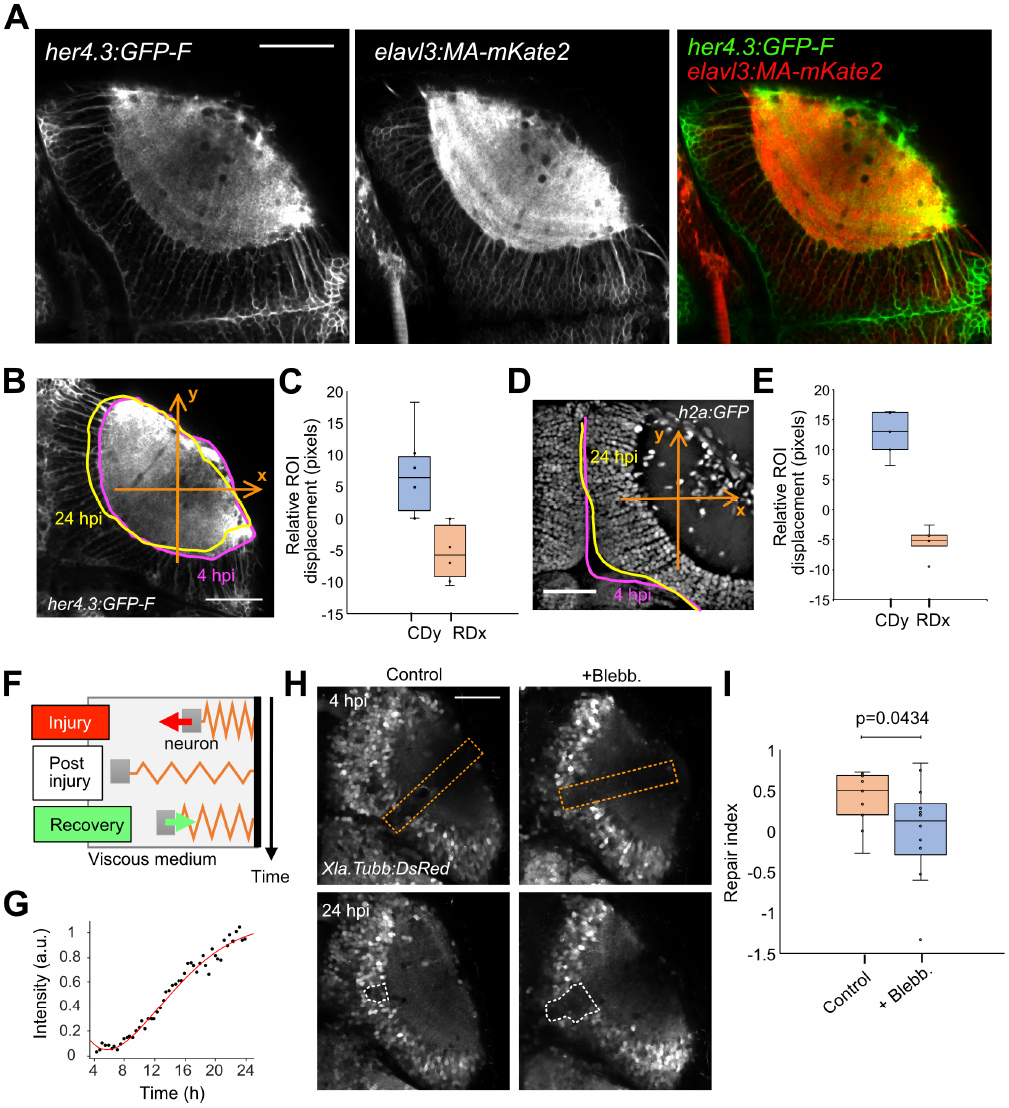
Optic tectum wounds close via tissue deformation. (A) Fluorescent images of a Tg(her4.3:GFP-F);Tg(elavl3:MA-mKate2) fish at 4 dpf. (B) Neuropil deformation measurement using Tg(her4.3:GFP-F);Tg(elavl3:MA-mKate2) larvae. Deformation is measured along the axes shown (orange arrows). (C) Quantification of deformation of the neuropil contour using Tg(her4.3:GFP-f) larvae (n = 6) and of the (E) PVZ delimitation Tg(h2a:GFP) larvae on the caudal (y-axis, CDy, blue) and rostral (x-axis, RDx, orange) axes (n = 6). (D) Deformation measurement using, with deformation measured along the axes shown (orange arrows). (F) Model of viscoelastic dynamics of neuron displacement comprising three phases; see Suppl. Information). (G) Example of experimental wound closure curve (black dots) fitted by the model (red curve). (H) Example images of injured optic tectum of control and blebbistatin treated Tg(Xla.Tubb:DsRed) animals at 7 and 24 hpi. The orange dashed lines show the position of the injury. The white dashed lines show the analysis regions for calculating the repair index. (I) Quantification of the repair index (RI) for control (n = 9) and blebbistatin-treated (n = 12) fish. T-test p value <0.05. Box plots show the median, box edges represent the 25th and 75th percentiles, and whiskers show the full data range. Scale bars represent 50 µm on all images. All quantifications are between 4 and 24 hpi unless otherwise stated.

Indeed, astrocytic processes support the structural integrity of the optic tectum and should be involved in any whole-tissue movement or deformation [15, 16]. We observed that the neuropil underwent deformation, with a contraction of the caudal part of the tectum and a dilation perpendicular to the curved rostrocaudal extent in the rostral part of the tectum (Fig. 2B,C). We measured the ventricular border deformation using Tg(h2a:GFP) larvae (Fig. 2D). The rostral part showed a deformation towards the contralateral side, and the caudal part showed a deformation towards the rostral part (Fig. 2E). This is consistent with the deformations observed for the neuropil and with the concerted trajectories of individual cells described above. Hence, the injured optic tectum tissue appears to be subject to anisotropic deformations during wound closure.

### Wound closure is due to mechanical forces that induce tissue deformations

Tissue deformation during wound closure could result from passive movement of the tissue into the gap or by active forces pulling on the tissue. To investigate whether the observed kinetics of wound closure could be explained by active forces, we developed a theoretical model that assumes elastic forces acting on the tissue. This model assumes that neurons are bound to a spring in a viscous medium (Fig. 2F) and follow the dynamics of a viscoelastic harmonic oscillator. The spring represents the elasticity of axons bundled to each other by protocadherins [17] and of astrocytic processes embedding neuron cell bodies (localised in the PVZ as shown by images of their nuclei) and axons. We considered only one half-period of oscillations as we observed the displacement to be monotonous towards the neuropil. We generated theoretical displacement curves of neurons, and, considering them as landmarks of the PVZ tissue, we compared them to the experimental wound closure kinetic curves. The theoretical curves obtained from the model were in close agreement with the experimental kinetic curves (Fig. 2G). This suggests that an external elastic force could drive neural tissue displacement.

To experimentally test whether active forces could be involved in tectum repair, we used blebbistatin, a drug that inhibits non-muscular myosin II activity and thus reduces force generation by cells relying on the actomyosin complex [18]. We first performed brain injuries using the same experimental conditions and then treated injured larvae with 1 µM blebbistatin from 7 to 24 hpi. This delayed drug application allowed microglia to migrate to the injury site [3]. To quantify the efficiency of wound closure and the effect of blebbistatin, we defined a repair index as the ratio of the injury volumes at 4 and 24 hpi. We used this approach on 9 control and 12 treated fish and found that wound closure was significantly impaired in treated larvae (Fig. 2H,I).

Altogether, these analyses suggest that, after injury of the optic tectum, the wound closes via tissue deformation caused by mechanical forces involving the actomyosin complex.

### Mechanical forces originate from a region where microglia accumulate

We next investigated where the mechanical forces responsible for the wound closure originated. Using cell nuclei as landmarks of the tectal PVZ to characterise local tissue deformations, we performed a displacement field analysis on data presented in Fig. 1. This showed that trajectories of nuclei converged towards the neuropil at an average normalised tectum radial position of 0.6, with an average angular position at 37º (Fig. 3A,B; see Suppl. Information - Viscoelastic model of neuron dynamics). This indicated a relatively narrow singular origin of potential contraction forces in the superficial neuropil centred over the injury site.

**Fig. 3.**
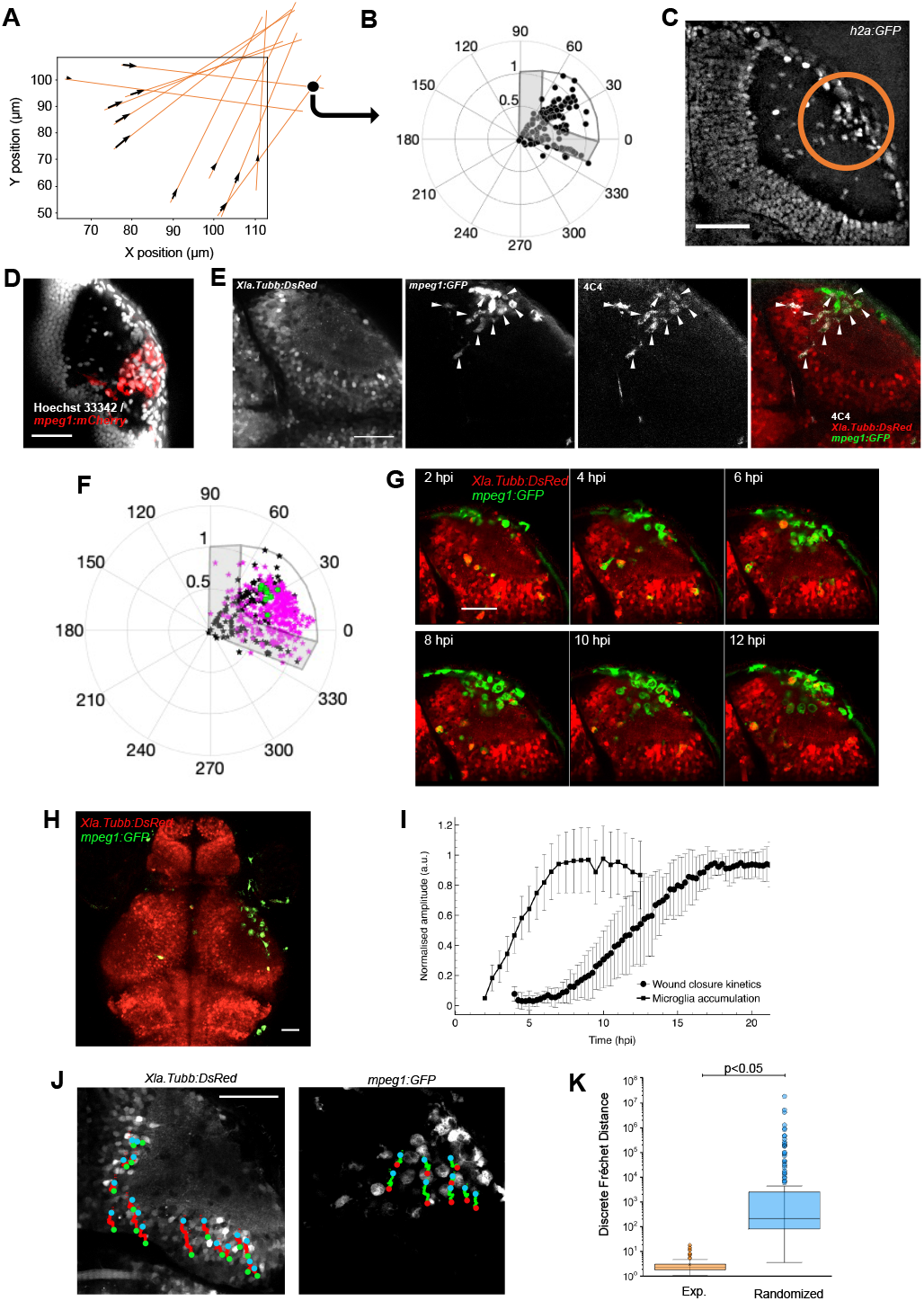
Microglia invasion in the neuropil correlates with the kinetics of wound closure. (A) Displacement field analysis showing the direction of displacement of individual neuronal nuclei (black arrows). The orange lines show how the intersection points (black spots) between trajectories are estimated. (B) Polar representation (in degrees) of the position of intersection points of individual trajectories of neuronal nuclei from 10 animals after tectum size normalisation and rotation. The optic tectum is represented as an overlay (boundaries in dark grey, PVZ in light grey). (C) Accumulation of cells in the neuropil (inside the orange circle) after injury in Tg(h2a:GFP) larvae at 20 hpi. (D) Identification of the accumulated cells as mpeg1+ cells using Tg(mpeg1:mCherry) (red) larvae, combined with nuclear staining (Hoechst 33342, grey). (E) Identification of the mpeg1:GFP+ cells as microglia using 4C4 immunofluorescence in an injured Tg(Xla.Tubb:DsRed);Tg(mpeg1:GFP) fish. White arrows point out mpeg1:GFP+/4C4+ double-labelled cells. (F) Polar representation of microglia positions (N=308 from 15 animals, magenta) in the neuropil compared to intersection points of trajectories of neuronal nuclei (black, green spots: centre of gravity of intersection points for each animal). (G) Selected frames of a time-lapse imaging series of an injured Tg(Xla.Tubb:DsRed);Tg(mpeg1:GFP) transgenic animal show the recruitment and accumulation of microglia in the injury site. (H) Lower magnification image of an injured Tg(Xla.Tubb:DsRed);Tg(mpeg1:GFP) transgenic animal showing that microglia did not accumulate in the intact contralateral tectum. (I) Kinetics of microglia accumulation from 2 hpi to 13 hpi (black square) and the average wound closure kinetics (black dots). (J) Dual-channel single cell tracking of neuronal cell bodies (Left, red trajectories) and microglia (right, green trajectories) in an injured Tg(Xla.Tubb:DsRed);Tg(mpeg1:GFP) larva. (K) Boxplot representation of the discrete Fréchet distance estimation between microglia trajectories and experimental (orange) and randomised (blue) trajectories of neuronal cell nuclei. Box plots show the median, box edges represent the 25th and 75th percentiles, and whiskers show indicate ± 1.5 x interquartile range. T-test p-value <0.05. Scale bars represent 50 µm on all images.

To characterise the origin site of the contraction force *in vivo*, we re-examined images of injured Tg(h2a:GFP) larvae. We identified an accumulation of cell nuclei near the convergence point of neuronal trajectories in the neuropil in all animals (n = 11; Fig. 3C). These cells could be microglia, based on previous observations that these cells are recruited in the neuropil of zebrafish larvae in the first 2 hours after injury [3]. Using Tg(mpeg1:mCherry) transgenic larvae, we observed that the accumulated cells were labelled by the transgene mpeg1, a marker of macrophages and microglia (Fig. 3D). We further identified the cells as microglia using immunofluorescence with the 4C4 antibody, which specifically recognises microglia [19, 20] (Fig. 3E). To confirm that, the accumulation site of microglial cells corresponds to the potential centre of force generation, we compared the position of the convergence points of the trajectories of neuronal cell nuclei, which we took as landmarks of the direction of tissue movement, with the position of microglial cells. We analysed the position of 308 individual microglial cells from 15 Tg(mpeg1:mCherry) larvae at 8 hpi, a time point when tissue movement after injury was fast (Fig. 3F, Fig. S3). We determined the accumulation centre to be in the neuropil at a normalised radial position of 0.88 and an average angular position of 17º. This is close to the centre of convergence of the neuronal trajectories determined earlier and is consistent with microglia being at the origin of the tissue pulling forces.

Next, we determined the temporal correlation between microglia accumulation and wound closure. We first performed time-lapse imaging on injured Tg(Xla.Tubb:DsRed);Tg(mpeg1:GFP) double transgenic larvae (Fig. 3G, Movie S2), in which neuronal cell bodies and microglia were fluorescently labelled in the same animal. This showed that microglia were recruited from 2 hpi (first time point of observation), rapidly accumulated until 6 hpi, and stayed accumulated in the neuropil for at least 12 hpi (Fig. 3G). Additional observations in histological preparations showed that microglia remained accumulated for at least 24 hpi (data not shown). Without injury, microglia did not accumulate and were rarely present in the neuropil itself in contralateral hemispheres or uninjured brains (Fig. 3H). Microglia move mainly in the periventricular zone between neuronal somata in these unlesioned situations. By comparing the kinetics of wound closure and microglia accumulation in the neuropil, we determined that the wound starts to close only after microglia have fully aggregated in the neuropil at 6 hpi (Fig. 3I). This is consistent with microglia as the origin of contraction forces.

To analyse the spatio-temporal correlation between microglia and tissue local deformations in more detail, we generated additional time-lapse movies of Tg(Xla.Tubb:DsRed);Tg(mpeg1:GFP) larvae at a higher frame rate to track microglia and neuronal cell bodies at the same time. We tracked neuronal cell bodies as tissue landmarks and microglia to compare their trajectories after injury. We observed that during the microglia accumulation phase (after 6 hpi), the tissue was deformed (measured by tracking neuron nuclei) towards the neuropil. Local deformations appeared to follow the movement of microglia compacting at the injury site (Fig. 3J). To estimate the similitude in terms of shape between trajectories, we computed the Discrete Fréchet distance between microglia trajectories and the experimentally observed neuronal trajectories. To estimate whether the value represents a good match between the movements, we compared it to that for randomised neuronal trajectories (Fig. 3K). The much higher Fréchet distance when using randomised trajectories indicated that local tissue deformation dynamics correlated well with microglia movement.

Overall, these results support the hypothesis that microglia accumulation is at the origin of the elastic forces inducing tissue deformation and, therefore, wound contraction.

### Computational modelling indicates sufficiency of microglia pulling behaviour for wound repair

During their migration microglial cells likely exert forces on their surroundings [21]. To estimate whether forces exerted by microglia could be sufficient to explain wound closure, we first took a theoretical approach by building a minimal system mimicking the brain tissue and microglia activity. To this aim, we developed a multi-agent system using the PhysiCell framework (Fig. 4A and Suppl. Information) using 3 agents: neurons, microglia, and skin cells. In the absence of external forces, neurons were allowed only to have a small random displacement as observed experimentally in intact fish. Skin cells were immobile and introduced to mimic the external tissue boundaries and stiffness.

**Fig. 4.**
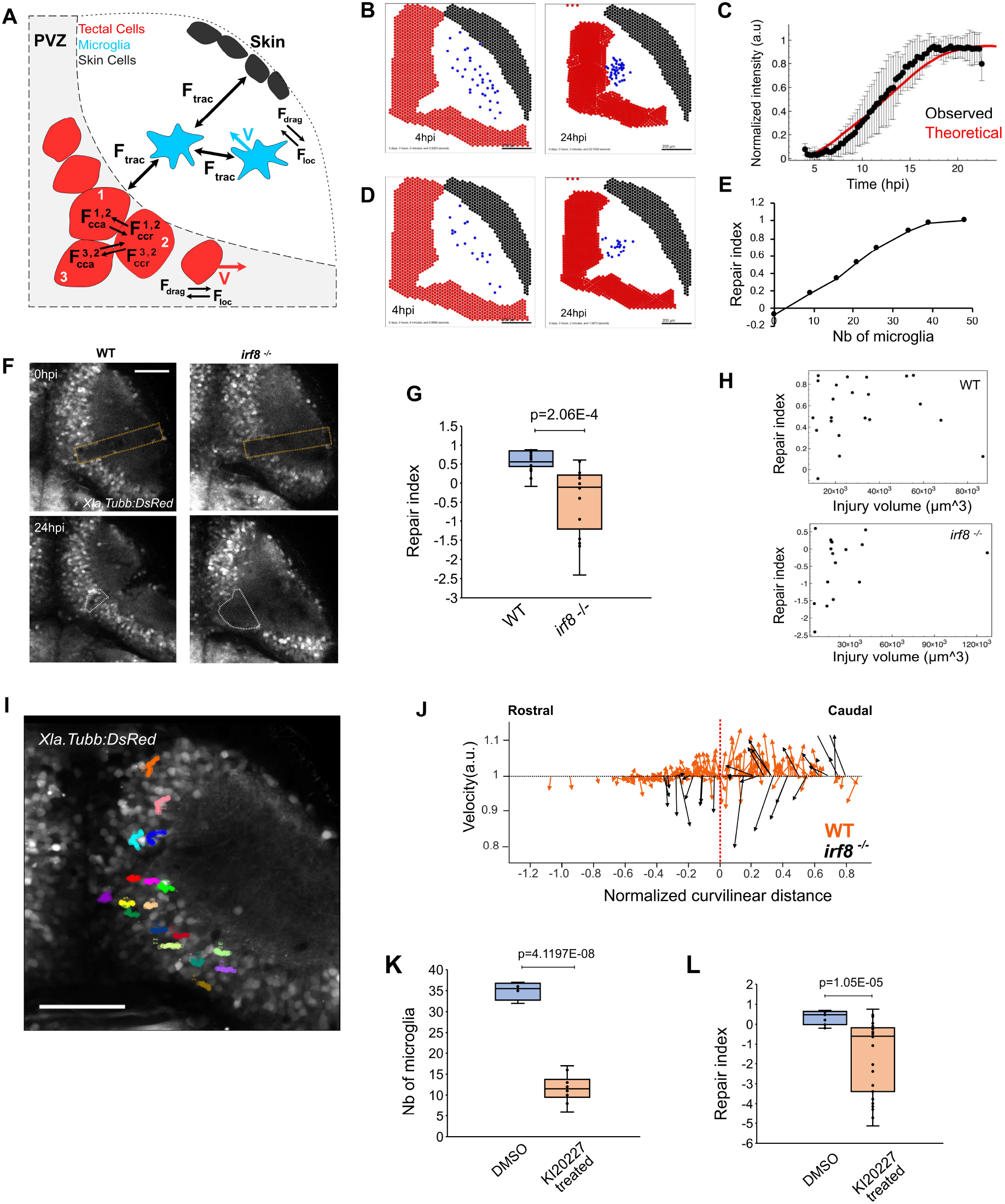
Microglia are necessary for brain tissue repair. (A) Graphic representation of the multi-agent model to simulate microglia-dependent wound closure. Blue: microglia, red: neurons, black: skin cells. The force vectors correspond to the forces described in the model (see Suppl. Information). (B) Example of simulation result showing microglia accumulation and wound closure. Left: simulated tissue after injury with patches of microglia agents in the neuropil. Right: the same simulation at 24 hpi. (C) Comparison of the kinetics of wound closure from *in silico* (red curve) and *in vivo* experiments (black dots, from Fig. 3I). Points: average value, bars: SD. (D) Simulation result with a low number of microglia compared to B, showing an impaired wound closure and lower degree of accumulation of microglia. (E) Relation between the repair index estimated on the simulated data and the number of microglia agents in the simulation. (F) Images of an injured tectum in wild type and irf8-/- Tg(Xla.Tubb:DsRed) larvae, right after the injury (0 hpi) and at 24 hpi. The orange dashed lines show where the injury was made. The white dashed lines show the analysis regions for calculating the repair index. Scale bar: 20 µm. (G) Quantification of the repair index (= 1 - V@24hpi / V@4hpi) in wild type (n = 21) and ifr8 mutant larvae (n =17). T-test p-value <0.001. (H) Correlation analysis of the repair index vs initial injury volume shows that the repair index is independent of the initial size of the injury. (I) Trajectories of neuronal cell bodies in an injured irf8 mutants (Scale bar: 50 µm). (J) Comparison of anisotropy of trajectories in wild-type (orange, from Fig. 1I) and in ifr8 mutants (black). (K) Quantification of the depletion of microglia using KI20227 T-test p-value <0.0001 (Controls n=4, treated n=10). (L) Measurement of the repair index in control (n=12) and KI20227 treated (n=26) animals. T-test p-value <0.0001. Box plots show the median, box edges represent the 25th and 75th percentiles, and whiskers show the full data range. Scale bars represent 50 µm on all images.

Microglia were allowed to migrate and exert elastic forces on all the other agents, following our first model of elastic forces controlling the displacement of neurons (see Fig. 3F). We then used this system to perform *in silico* experiments with an initial microglia distribution as isolated cells spread in the neuropil. We could reproduce both microglia accumulation and wound closure due to tissue contraction (Fig. 4B, Movie S3). Moreover, the kinetics of wound closure measured by the intensity of neuronal cells in the injury site over time (similar to the method used for experimental data) presented a curve that was a good fit for the experimentally observed wound closure curve (Fig. 4C). When microglial cells were reduced in number or even absent in the simulation (Fig. 4D,E), we found a linear relationship between wound closure, expressed as repair index, and microglia number until a plateau is reached when the number of microglia is sufficient to induce a full closure. In the extreme case where no microglia were present, the wound did not close. Therefore, our model predicts that the repair process depends on the number of microglia accumulating and that mechanical forces exerted by these cells may be necessary and sufficient to induce brain tissue contraction.

### Microglia are essential to restore the optic tectum architecture after injury

To experimentally test the model’s predictions, we first depleted microglia genetically, using the irf8 mutant [22], and pharmacologically, using the Csf1r inhibitor KI20227 [23]. Irf8 is a transcription factor that is essential for the development of all macrophages, including microglia. Irf8 mutants are adult-viable but selectively lack microglia and macrophages [22]. To visualise the tectal tissue the larvae also carried the Tg(Xla.Tubb:DsRed) transgene, which labels neuronal tissue. We observed that in 16 out of 21 wild-type control animals, the wound was closed at 24 hpi, as before (R.I. > 0.5). In contrast, the wound was closed in only 2 of 17 irf8 mutants (Fig. 4F).

We then compared the repair index between wild-type and irf8 mutants at 24 hpi (Fig. 4G), indicating a significantly lower repair index in the mutant. This was independent of the initial injury volume because there was no correlation between the initial injury volume and repair index across all animals (Fig. 4H). We then measured the trajectories of neuronal cell bodies in irf8 mutants to assess the tissue deformation. We found that trajectories were directed more often towards the outside of the tectum, with a larger amplitude for those oriented opposites to the neuropil, compared to injured wild-type larvae (Fig. 4F,G). Furthermore, from these measurements, we determined that the injury volume increases in Tg(Xla.Tubb:DsRed);Tg(irf8-/-) larvae in 59 % (10 of 17 animals) of the cases and never in wild type larvae. In these mutant animals, the wound enlarged, and the tectum dilated instead of contracting. Similarly, the Ki20227 treatment, which markedly depleted microglia (Fig. 4H), reduced the repair index in treated animals (Fig. 4I). Hence, our manipulations confirm our model’s prediction that in the absence of microglial cells, tectal wounds are not repaired. This indicates that microglia are necessary for wound closure.

### Microglia interact with astrocytic processes

To directly observe how microglia interact with the surrounding tissue during accumulation, we focussed on astrocytic processes because these are finely branched and widely distributed throughout the tectum. To visualise these astrocytic processes in fine detail, we used our newly generated Tg(her4.3:GFP-F) line. We performed 3D time-lapse image acquisitions on injured Tg(her4.3:GFP-F);Tg(mpeg1:mCherry) double-transgenic larvae, in which astrocytic processes Tg(her4.3:GFP-F) and microglial cells Tg(mpeg1:mCherry) were fluorescently labeled in the same animals, during a 4-24 hpi time window.

We observed an increase in the fluorescence signal for astrocytic processes in maximum intensity projections around the microglia accumulation site (Fig. S6C, Movie S4), which may indicate interactions between astrocytic processes and microglial cells. We then assessed at high magnification if the alteration of the meshwork of astrocytic processes was related to microglia dynamics by performing fast time-lapse imaging on injured Tg(her4.3:GFP-F);Tg(mpeg1:mCherry) larvae, starting at 6 hpi, when microglia were stably present. We observed astrocyte and microglia interactions (Fig. 5A, Movie S5) that seem to typically occur in three consecutive steps that we termed SAT: 1. S - sensing 2. A - adhesion 3. T – traction In phase 1 (sensing), microglia generate transient protrusions in apparently random directions and occasionally form longer-lasting associations with astrocytes shown by the colocalisation of both signals (phase 2, adhesion). During phase 3 (traction), microglia retract their protrusion leading to the deformation of astrocytic processes following the retracting microglia process. We observed more generally that astrocytic processes seemed to be pulled by microglia when microglia moved (“migration” events) inducing additional deformation of the meshwork (Fig. 5A,B; Movie S6).

**Fig. 5.**
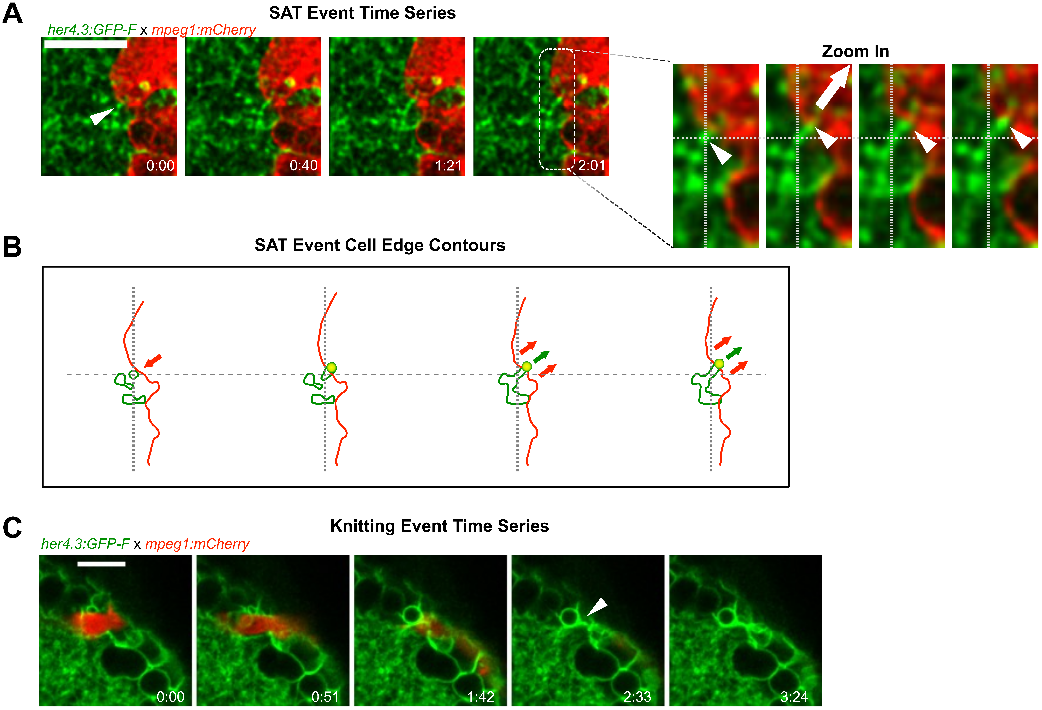
Microglia contacts displace astrocytic processes. (A) Fast time-lapse imaging sequence on a Tg(her4.3:GFP-F);Tg(mpeg1:mCherry) larva showing an SAT process. The white arrow points out the adhesion event. (A’) Cropped sequence from A. The small white arrows point to astrocytic protrusion pulled by microglia protrusion. The big arrow shows the direction of the traction force. The dashed line indicates the initial and final positions. (B) Outline of microglia (red) and astrocytic structures (green) from images in e. Arrows show the direction of displacement. (C) Example of a knitting event. All scale bars: 10 µm.

More surprisingly, we observed an original phenomenon that we named “astrocytic knitting” where astrocytic processes seem to be “glued” together after the traction phase (Fig. 5C, Movie S7). Overall, from the series of 30-40 min timelapses acquired at 6 hpi, we observed 54 SAT events from the analysis of 43 microglial cells accumulated in the neuropil (from 3 animals) and estimated an average frequency of 20 SAT events/hour and per hemi-tectum. The frequent occurrence of the SAT processes during the repair phase shows that microglia transiently attach to astroglial processes during their migration, leading to their deformation. This interaction could lead to tissue contraction.

These observations are consistent with the notion that microglia exert a mechanical action on the tissue, by pulling on astrocytic processes.

### Micro laser ablation of microglia-astrocytic contacts leads to retraction of processes

To obtain direct evidence for a microglial pulling action on astrocytic processes, we targeted these contacts in wild-type animals with a 2-photon laser and imaged a 3D ROI within the injury site in Tg(her4.3:GFP-F);Tg(mpeg1:mCherry) double-transgenic larvae at 16-18 hpi at every 15 seconds for 5 minutes. We measured the change in area of microglia and astrocytic processes. For the microglia, we observed a large retraction of the mostly elongated contact site within the first frame (6/6 cases; Fig. 6A,B, S8A; movie S9).

**Fig. 6.**
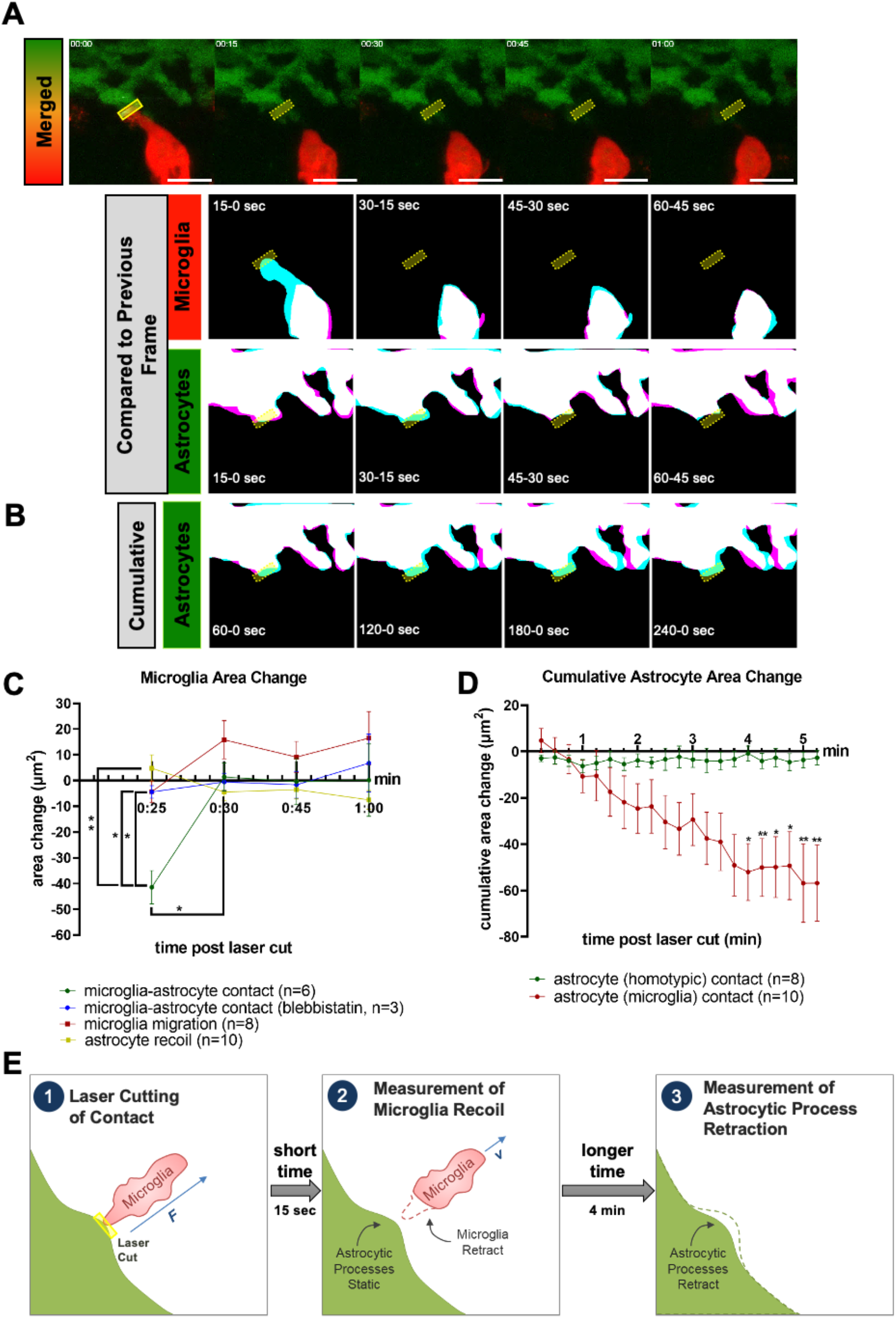
Microglia and astrocytic process recoil following laser severance of contact sites. (A) Diagram of laser-induced microglia recoil away from astrocytic contacts and slower retraction of astrocytic processes. (B) Time-lapse images of microglia (red) and astrocytes (green) following laser ablation (upper, up to 1 min post-ablation), with area masks overlaid between adjacent time points for microglia (middle) and astrocytes (lower). (C) Cumulative area masks of astrocytes for 1-4 min post-ablation. (D) Change in area following laser ablation for microglia which were contacting astrocytes (green, n=6), blebbistatin treated microglia contacting astrocytes (blue, n=3), microglia migration (red, n=8), and astrocytes which were contacting microglia (yellow, n=10). Area change represents the difference between T(n+1) - Tn as shown in (B) as magenta and blue, respectively. (E) Cumulative area change for astrocytes with homotypic (green, n=8) or microglia (red, n=6) contacts after ablation. Area change represents the total area changed between T0 and the given time point. Statistics were assessed by 2-way ANOVA with Šídák’s multiple comparisons test; P < 0.05 = *, P<0.01 = **. All scale bars represent 10 µm, and all ablation sites are marked with yellow boxes on montages.

Quantifying the area change of the microglia indicated a change in the first 15 seconds after injury that was 30-fold larger than in any subsequent frame, and more than 3-fold greater than in microglia migrating towards a laser-injury site from a distance (Fig. 6C, Fig. S4B). Of note, microglia with severed processes resumed movement after 30-45 seconds (6 out of 6 cases) and, hence, were not destroyed by the laser.

Since we observed impairment of wound closure in animals treated with blebbistatin (Fig. 2I), we repeated the experiments using this treatment. As expected, the initial area change was strongly reduced compared to experiments in which microglia-astrocytic contacts were severed in the absence of blebbistatin (3/3 cases Fig. 6C, Fig. S4C). To achieve higher temporal resolution, we recorded only one optical section with a time resolution of 0.65 seconds. We managed to observe two thin microglial processes that retracted almost immediately. Within the first 3 seconds after severing the contact (Fig. S5, movie S10) they moved at an average speed of 52 µm/min (0.89 µm/s). After that, processes move towards the injury site again at a much lower speed (0.18 µm/s), indicating survival of the cells and migration towards the injury. Hence, also at higher temporal resolution, recoil speed greatly exceeds the migration speed of microglia. This indicates that microglial processes after laser severance quickly retract, consistent with the contact being under tension.

Astrocytic processes showed retraction in the opposite direction to the microglia (Fig. 6A-E). However, within the first minute after severing the contact, we did not observe significant movement of the astrocytic processes (Fig. 6A,C).

Analysing the change in area for the astrocytic processes for a longer time period indicated that their reaction was more protracted and cumulative area change became significant only at 4 min after cutting off the contact with microglia (Fig. 6B,D). Injuring astrocytic processes in places without microglia in the vicinity (> 5 µm) did not lead to a displacement, indicating dependency of the movement on previous microglia contact (Fig. S4D). Hence, astrocytic processes show a specific, but slow reaction to the severance of microglia contacts. Retraction of astrocytic processes and microglial cells from a severed contact point provides direct evidence for a mechanical interaction between these cell types (Fig. 6E).

In summary, our micro laser ablation results support the specific mechanical role of microglia in wound closure after brain injuries.

### Microglia action on wound closure depends on lcp1 gene function

If microglial cells are responsible for tissue deformation by exerting local traction forces on the surrounding tissue, they need to contain non-muscular myosin II in addition to actin. We confirm this here by immunofluorescence against non-muscle myosin IIB/myh10 in Tg(mpeg1:GFP) animals, showing enrichment in tectal microglial cells (Fig. S6B). To detect the actin cytoskeleton, we used Tg(mpeg1:GFP);Tg(beta-actin:utrophin-mCherry) animals. These are double-transgenic animals, in which the actin-binding protein utrophin is transgenically labelled with mCherry and microglial cells in the tectum are labelled by GFP. In these animals, we observed strong labelling of the microglial cell cortex with mCherry, compared to the cell’s center and the surrounding tissue in the neuropil (Fig. S6A). Moreover, we observed a close association of the actin cytoskeleton in microglia with SAT events (Fig. S7; movie S8). This suggests the presence of the cellular machinery for force generation in microglia during tissue deformation.

To determine whether the magnitude of microglia traction force is relevant to wound closure mechanics, we decided to weaken this force in a cell type-specific manner in microglia. To achieve this, we decided to ablate L-plastin function in microglia. L-plastin is coded for by the gene lcp1, which is specifically expressed in immune cells under physiological conditions. In our tectum lesion assay, only microglia are present (see 4C4 immunofluorescence in Fig. 3E and [3]. L-plastin stabilises bundles of filamentous actin (F-actin) and is thus essential for force generation, for example in cancer cell invadopodia [24]. Importantly, F-actin is also highly enriched in protrusions of brain macrophages that exert traction forces on endothelial cells during blood vessel repair [25], suggesting that lack of L-plastin weakens F-actin function in traction events.

First, we confirmed the expression of lcp1 in microglia. To do so, we performed double-HCR for the microglia marker apoeb [26] and lcp1 and observed accumulation of lcp1 expressing cells in the injured tectum at 6 hpi (Fig. 7A). Of these cells, 92 *±* 3.69 % were apoeb positive, indicating nearly exclusive expression of lcp1 in microglial cells in the injured tectum (Fig. 7B). Hence, any manipulation of lcp1 will be specific to microglia cells.

**Fig. 7.**
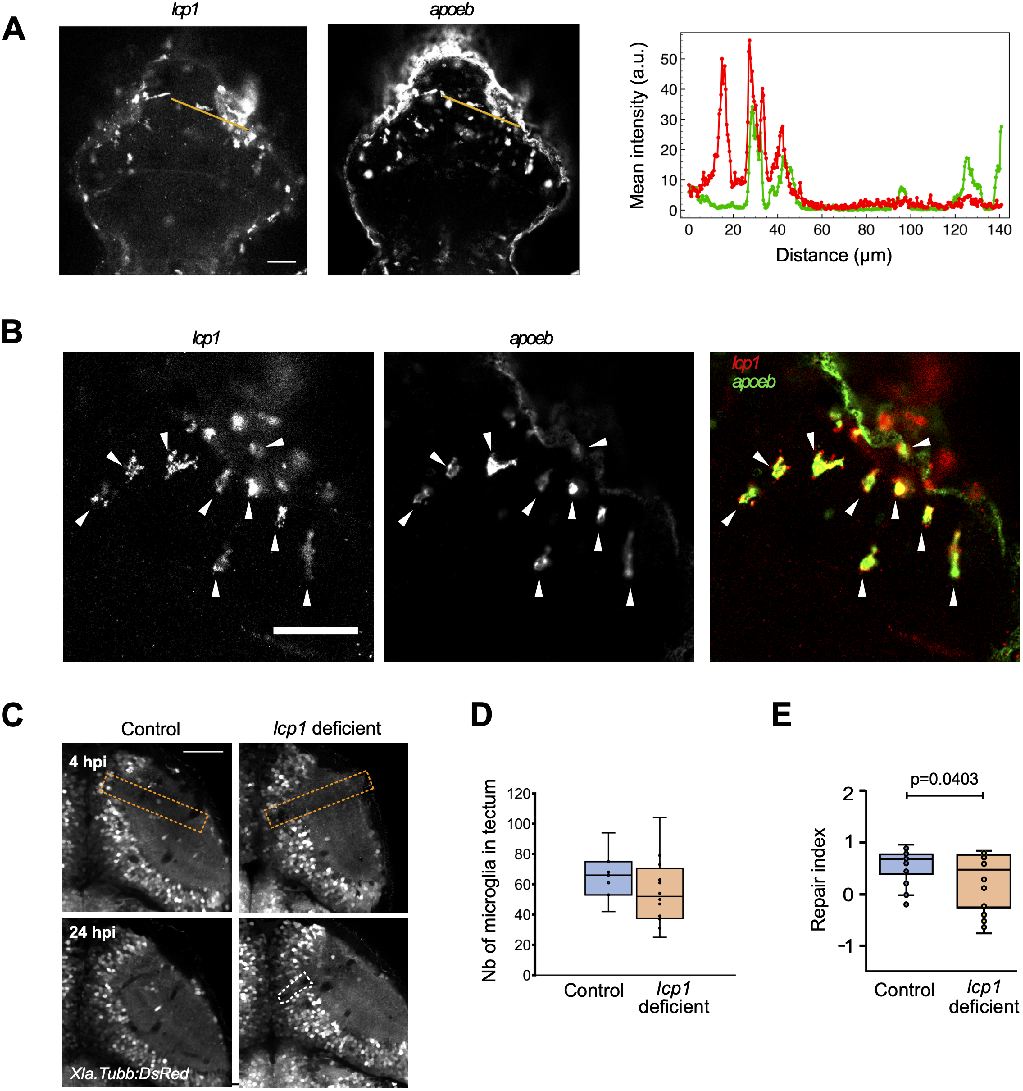
Microglia action relies on lcp1. (A) Expression of lcp1 and apoeb as detected by hybridisation chain reaction fluorescent *in situ* hybridisation (HCR FISH) in injured larvae at 6 hpi. The intensity profiles shown on the graph (red: lcp1, green: apoeb) were measured along the orange lines shown in the images. (B) Zoomed in HCR FISH images for lcp1a (green) and apoeb (red), showing an almost complete overlap of patterns. Arrows point to lcp1+/apoeb+ cells at the lesion site (C) Representative images of wound closure in control and haCR-injected animals are shown; orange dashed lines: injury site, white dashed lines: wound delimitation in the PVZ. (D) Boxplot representation of the repair index in all haCR-injected animals. (p-value from unpaired t-test). (E) Boxplot representation of the number of microglia in the optic tectum at 24 hpi in haCR-injected animals. All t-tests were non-significant (p>0.05). For all experiments, n = 4 for control injected, and n = 10 for haCR-injected animals. All scale bars represent 50 µm.

To functionally analyse the relevance of lcp1 in our wound closure assay, we designed highly active sCrRNAs (haCRs) to effectively create somatic mutants [27]. Restriction Fragment Length Polymorphism (RFLP) analysis after zygote injection confirmed that haCRs were indeed highly efficient *in vivo* in disrupting a targeted restriction enzyme recognition sequence (Fig. S8). Somatic mutation of lcp1 did not overtly impair the accumulation of microglia in the injury site and no differences in the number of microglial cells were observed (Fig. 7C), indicating that microglia could still exert potential functions that were unrelated to force generation. However, in lcp1 somatic mutants, the tectum wound did not close completely and quantification indicated that the repair index was significantly lower than in controls (Fig. 7D, E). Hence, the F-actin bundling lcp1 in microglia is necessary for effective wound closure by microglia.

These observations indicate that forces exerted by microglia make an essential contribution to brain wound closure.

## Discussion

Here we present evidence that microglia play a previously unknown role in the central nervous system (CNS) tissue repair by facilitating contraction of the injured tissue, leading to the closure of a physical wound.

Direct evidence for the role of microglia in pulling on surrounding tissue comes from our observation that microglia displace astrocytic processes and that severing microglia/astrocyte contacts leads to a rapid pull-back of microglia and a protracted pull-back of astrocytes. This indicates physical tension between these cellular elements. At high temporal resolution, we observed an almost immediate withdrawal of microglial processes at speeds that were consistent with other paradigms testing tissue tension [28, 29, 30, 31]. The reasons for the slower, but consistent movement of the astrocytic processes need further investigation but may lie in their meshwork-like arrangement, rather than being isolated entities, like the microglial cells. Our observations are supported by other studies showing that microglia can exert traction forces on their substrate and sense differences in tissue stiffness [10, 32]. In addition, the role of macrophages in blood vessel repair with direct mechanical action at the cellular scale has been demonstrated *in vivo* [25]. Recently, a study has demonstrated the importance of microglia for the maintenance of the mechanical properties of brain tissue in zebrafish larvae [33].

We observed directed tissue movements and microglia accumulation correlated with wound closure. This prompted a modelling-based approach that predicted that mechanical forces by microglia could be sufficient for wound closure. Our mathematical model is highly simplified, for example, assuming equal forces between microglia and other cellular elements. Nevertheless, even with these minimal assumptions the model accurately predicted impaired wound contraction in the absence of microglia in irf8 mutants and reduced abundance after Csf1ra inhibition and thus suggests that microglia pulling action may be sufficient to close a brain wound in our model. However, the absence of microglia is likely to also impact the outcome of injuries indirectly. For example, the lack of cytokine signalling to astrocytes [11] or other cell types is likely to contribute to impaired wound closure. In spinal cord regeneration, macrophage-derived cytokines are crucial [34]. Nevertheless, in experiments targeting cytoskeleton function, i.e. blebbistatin administration, as well as in lcp1 deficient animals, microglia were still present in the tectum, suggesting that at least some of their signalling functions were retained. Therefore, these experiments lend support to the hypothesis that microglia have a role in tissue force generation that is essential in tissue wound closure.

Regarding the cellular machinery involved in the pulling action of microglia, we find that microglia-specific lcp1 is necessary for efficient wound closure of the tectum. The protein product L-plastin stabilises F-actin bundles that are critical for force generation in stress fibres [24]. Lcp1-deficient microglia are the only lcp1-expressing cell type in the injured tectum. Hence, the loss of lcp1 in microglial in lcp1 somatic mutants specifically weakens F-actin-dependent force generation in microglia. Ablation of lcp1 function thus supports that the tension between astrocytic processes and microglia is sufficiently large to make a meaningful contribution to wound closure. Interestingly, lcp1-deficient macrophages undergo shape changes after tail injury in larval zebrafish, consistent with an inability to exert pulling forces [35].

Failure to close the wound in the absence of microglia could also be explained by the lack of debris removal, another microglial function [3]. However, in injured irf8 mutants, neutrophils invade the tectum, which they did not do in wild-type animals, and were able to clear dead cells when microglia were missing [8]. This suggests that the inability to close a wound, at least in the irf8 mutant, is unlikely to be a consequence of impaired removal of dead cells.

The role of microglia in human TBI is not fully understood. Increased microglia inflammation leads to secondary damage [36, 37], but may also have positive functions for recovery [38]. Human microglia and zebrafish telencephalic microglia show similar gene expression signatures after injury [39]. Interestingly, granulin-dependent clearance of lipid droplets and TDP-43 may be an important difference in the inflammation status of microglia between the species [39]. Our results point to a perhaps even earlier role of microglia in mechanically restoring tissue integrity and thus promoting healing of the brain tissue.

The accumulation of microglia has also been observed in other CNS injury contexts, for example in stroke [38] and spinal cord injuries in mice [40, 41], suggesting wider relevance of our findings. Similar to our observations, microglia and/or macrophage accumulations are progressively surrounded by a dense meshwork of astrocytic processes in these models, in a process termed corralling [42]. Loss of microglia or microglia/macrophage-specific gene knockout of adhesion molecules, such plexin-b2, leads to disorganised and increased wound size [38, 40]. Conversely, inactivating the signalling molecule Stat3 in astrocytes leads to similar phenotypes, suggesting interactions between astrocytes and microglia [42]. The exact role of microglia in wound compaction or corralling is unclear in these injury models. Here, we have established an injury system in zebrafish where corralling-like cellular behaviour can be directly observed and manipulated.

In summary, we propose a brain repair mechanism in which mechanical forces in the early stages of tissue regeneration are crucial to contracting the tissue and thus facilitate subsequent repair processes. This physical aspect of tissue repair is rarely investigated and offers a more holistic view of the processes involved. Furthermore, our findings reveal a new role in the microglial response after an injury. These cells are indispensable for the mechanical action to close the wound after an injury.

Ultimately, our results may inspire therapeutic approaches targeting mechanical forces related to microglia and other cell types for healing traumatic brain injuries and other tissue damage.

## Materials and Methods

### Fish husbandry

All zebrafish lines were kept and raised under standard conditions [32] and all experiments were approved by the UK Home Office (project license no.: PP8160052) or according to German animal welfare regulations with the permission of the Free State of Saxony (project license no.: TVV36/2021). Following the guidelines of the 3Rs, we only used larvae aged up to 5 dpf. For experimental analyses, we used larvae of either sex of the following available zebrafish lines: Tg(Xla.Tubb:DsRed)^zf148^ [25]; Tg(betaactin:utrophin-mCherry)^e119^ [34]; Tg(h2a.F/Z:GFP)^kca6^ [43] (referred to as Tg(h2a:GFP)); Tg(mpeg1.1:GFP)^gl22^ [44]; Tg(mpeg1.1:mCherry)^gl23^ [44]; Tg(irf8)^st95^ [45]; Tg(elavl3:MA-mKate2)^mps1^ [46]. The Tg(her4.3:GFP-F)^mps9^ transgenic zebrafish line was established using the DNA constructs and methodology described below. If necessary, larvae were treated with 100 µM N-phenylthiourea (PTU) to inhibit melanogenesis. All chemicals were supplied by Sigma unless otherwise stated.

### Generation of Tg(her4.3:GFP-F) transgenic fish

To create the donor plasmid for generation of her4.3:GFP-F transgenic zebrafish, the sequence coding for the membrane-localised GFP (EGFP fused to farnesylation signal from c-HA-Ras) was amplified from the pEGFP-F vector (Clonetech) using oligos 5’-TTATTTATCGATCCACCATGGTGAGCAAGGGC-3’ and 5’-TTTATTATCGATTCAGGAGAGCACACACTTGCAGCT-3’ and cloned downstream of the her4.3 (previously known as her4.1) zebrafish promoter [36]. Transgenic fish were established by injection of 40 pg of the donor plasmid together with mRNA of the Tol2 transposase into 1-cell embryos [37].

### gRNA injections

The gRNAs were injected into the yolk at the one-cell stage of development. The injection mix was prepared on the morning of injections. The mix consisted of 1 µL Cas9 protein (BioLabs, M0369M), 1 µL Fast Green FCF dye (Sigma, 2353-45-9), 1 µL 250 ng/µL SygRNA-tracr (Sigma, TRACR-RNA05N), 1 µL gRNA and 1 µL nuclease-free water. When two gRNAs were co-injected, the nuclease-free water was substituted with the second gRNA. After mixing gRNAs and tracr (and water if using), the mixture was heated to 95 degrees for five minutes and then kept on ice for 20 minutes. After this, the Cas9 and dye are added, and the mixture is again heated to 27 degrees for 10 minutes. For every experiment two injection mixtures were made, one with the gRNA of interest and one with a control gRNA (5’-TTACCTCAGTTACAATTTAT-3’). lcp1 was targeted with a gRNA (5’-GAACCCGGUACCCCGGCAGA-3’) as previously published [19].

### RFLP analysis of gRNA efficiency

To test the efficacy of gRNAs targeting lcp1, we used the Restriction Fragment Length Polymorphism (RFLP) method as described in Keatinge et al. [27]. See the table below for primer sequences and corresponding restriction enzymes.

**Table 1.**
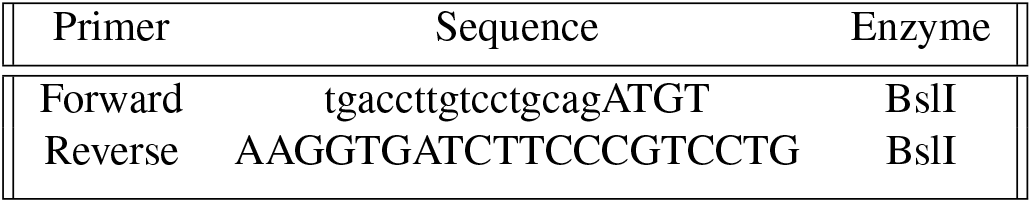
Primer sequences for lcp1.

Genomic DNA was extracted from individual embryos (1-5 dpf) at by heating the whole embryo with 100 µl of 50 mM NaOH at 95°C for 10 min followed by the addition of 10 ul of 1 M HCl. PCR products were generated using BioMix™ Red (Bioline, Bio-25006) and were subsequently incubated with the respective restriction enzyme (BioLabs) for 1.5 h at the specified temperature, followed by separation on a 2 % agarose gel (100 V for 40 min) and visualisation on a transilluminator.

### Hybridisation Chain Reaction Fluorescent *in situ* hybridisation (HCR FISH)

For HCR FISH, HCR probe sets were ordered from Molecular Instruments, targeting the following mRNAs: apoeb1 (NCBI Reference NM _ 131098.2) and lcp1 (NCBI Reference NM_ 131320.3). All HCR buffers as well as the h1 and h2 hairpins were purchased from Molecular Instruments; Los Angeles, California, USA. HCR FISH was performed as previously described in Choi et al. [47]. Larvae were fixed at 6 hpi in 4% PFA at RT for 3 hours. Fixed larvae were dehydrated in increasing concentrations of methanol and stored at -20°C in methanol overnight. Larvae were rehydrated in decreasing concentrations of methanol and washed 3 times in 1x PBST. Permeabilisation was performed by incubating the larvae for 40 min in 30 µg/ml Proteinase K (in 1x PBST). Larvae were post-fixed in 4% PFA at RT for 15 min. and subsequently washed 4 times in 1x PBST. Pre-hybridisation was performed with pre-warmed HCR hybridisation buffer (Molecular Instruments) for 30 min. at 37° C. 2 pmol of the respective HCR probes were diluted in 500 µl HCR hybridisation buffer. Hybridisation buffer was replaced with the HCR probe mix and hybridization was performed for 16h at 37°C. To remove excess probe, the larvae were first washed 4 times for 15 min. each in HCR Wash Buffer (Molecular Instruments) at 37° C, followed by several washes in 5X saline-sodium citrate buffer with 0.1% Tween20 (SSCT) at RT. Subsequently, larvae were incubated in HCR Amplification Buffer (Molecular Instruments) for 30 min at RT. Meanwhile, 30 pmol of respective hairpin h1 and h2 (Molecular Instruments) were prepared by snap-cooling 3 µl of the respective 10 µM hairpin stock. Snap-cooling was performed by incubating the hairpins for 90 sec. at 95° C and cooling them down for 30 min. at RT in the dark. Hairpin solution was prepared by transferring 10µl of the snap-cooled h1 and h2 hairpin to 500 µl Amplification Buffer. The amplification step was performed by incubating the larvae for 16 h in hairpin solution. Excess hairpin solution was removed by washing the larvae 4 times in 5x SSCT for 15 min each at RT. Larvae were stored in 70 % glycerol (in 1x PBST) at 4°C.

### Induction of brain injury

For inducing brain injuries, Zebrafish larvae at 4 dpf were anaesthetised using 0.016 % ethyl 3-aminobenzoate methane sulfonate (MS-222, Sigma). They were then mounted in 1% low melting point agarose (LMP, Life Technologies) with their dorsal side facing upward and their anteroposterior axis as horizontal as possible. The optic tectum was then injured under visual guidance via a stereomicroscope using a metal insect pin with a diameter of 80 µm (Fine Science Tools) mounted on a micromanipulator (Narishige or WPI). The tip of the metal pin was slightly bevelled to facilitate penetration of the skin, and the angle between the pin and the horizontal plane was 20-30°. For induction of injury, the tip of the metal pin was inserted into the optic tectum to a depth of 200 µm and immediately retracted. The injury procedure itself took less than 2 sec. After an injury, larvae were carefully released from the agarose and allowed to recover in fresh fish water for varying amounts of time depending on experimental requirements (typically 4 hours before imaging).

### Blebbistatin treatments

Animals were incubated with 1 µM para-nitroblebbistatin (Axol Biosciences or BIOZOL), referred to as blebbistatin, solubilised in double distilled H2O. The drug was added directly to the fish water 7h after the injury, and incubation continued until the time of experimental readout. The repair index was determined as described in the dedicated section.

### KI20227 treatments

KI20227 (Tocris) powder was dissolved in sterile filtered DMSO to produce a 50 mM stock. 2 dpf Tg(Xla.Tubb:DsRed);Tg(mpeg1:GFP) larvae were screened for transgenes, anaesthetised and transferred into wells with 500 µL of a 25 µM solution of KI20227 diluted in fish water with PTU. Larvae were incubated for 48h before injury or cell counting to verify the efficient suppression of microglia in the optic tectum (Fig. 4H).

### Immunofluorescence

All incubations were performed at room temperature unless stated otherwise. At the time point of interest, larvae were fixed in 4% PFA-PBS containing 1 % DMSO at 4°C overnight. Larvae were then washed in PBTx (1% Triton X100 in PBS). After permeabilisation by incubation in PBTx containing 10 µg/ml Proteinase K for 15 min, larvae were washed twice in PBTx. Then larvae were incubated for 1 hour in 4 % BSA-PBTx (blocking buffer) and incubated with primary antibody (mouse anti-HuC or donkey anti-Myh10) diluted at 1:100 in blocking buffer at 4°C overnight. On the following day, larvae were washed 6 times in PBTx for 20 min each, followed by incubation with secondary antibody (anti-mouse Cy3, Jackson) diluted at 1:300 in blocking buffer at 4°C overnight. The next day, larvae were washed 6 times in PBTx for 30 min each and twice in PBS for 15 min each, before mounting in 1% low-melting point agarose for imaging.

### HuC/EdU staining

Brain injuries were carried out on 4 dpf Tg(h2a:GFP) larvae as described above. After performing the lesions, larvae were immediately placed in a solution of 50 µm 5-ethynyl-2’-deoxyuridine (EdU; Sigma-Aldrich) and were allowed to develop in this solution.

At 5 dpf (24 hpi) the larvae were culled with an overdose of MS-222 and fixed with 4% paraformaldehyde (PFA) overnight on a rocker. Larvae were washed 3 times with PBTx and incubated in methanol for 5 min before being placed in fresh methanol at -20° C overnight. Samples were rehydrated in a dilution series (75, 50, 25%) from methanol to PBTx and then washed in PBTx. Larvae were digested using Proteinase K (10 µg/ml in PBTx) for 45 minutes at room temperature. They were then fixated for 15 min in 4% PFA at room temperature. After 3 washes in PBTx, the animals were incubated in a 1% DMSO/0.5% Triton X100 solution for 20 min and washed in PBTx. The EdU Click-iT reaction solution (Roche) was prepared according to manufacturer’s instructions and larvae were incubated at room temperature in the dark for 2 hours. All further incubations, including the immunofluorescence, were carried out in the dark. Larvae were thoroughly washed several times in PBTx before proceeding to immunofluorescence with anti-HuC to identify neurons.

### Image acquisition

Before live confocal imaging, zebrafish larvae were anaesthetised using 0.008% MS-222 and mounted in 1 % low melting point agarose (LMPA), with their dorsal side facing upward to image the optic tectum. The agarose was then covered in fish water with 0.008% MS-222 to prevent desiccation during imaging and animal movements.

For long-term imaging, the larvae were mounted in 1% low melting point agarose in a 90 mm Petri dish and placed on the microscope stage using a custom 3D printing adapter. The animals were maintained at a stable temperature of 28º C using a stage incubator with a bucket of water to prevent excessive evaporation of the fish water from the Petri dish. For z-stacks of the optic tectum in living or fixed samples, images were acquired with typically 2 µm intervals between optical planes. In injured animals, z-stacks were chosen such as to encompass the entire injury site.

For acquiring short time-lapses on living larvae or 3D stack on fixed animals, larvae were mounted in LMPA on a microscope coverslip. A reservoir was obtained by setting the coverslip on a microscope slide with a spacer made of vacuum grease and filled with fish water to keep animals hydrated.

All fluorescence images except for HCR FISH and laser cutting samples were acquired on a Zeiss LSM880 confocal microscope equipped with an Airyscan detector and a 20x NA 1.0 LWD water dipping objective lens.

Fluorescently HCR FISH stained larvae were recorded as confocal stacks. Confocal imaging was performed using a Carl Zeiss LSM 980 inverse laser scanning microscope. A 20x Plan-Apochromat, Air, DIC objective (Zeiss, NA=0.8) was used. The 561 nm and the 633 nm laser lines were used for confocal microscopy. The pinhole size was adjusted to 1 Airy Unit.

### Repair efficiency assay

The larvae were injured and imaged as described. After imaging at 24 hpi, the larvae were culled and genotyped individually by analysis of the restriction fragment length polymorphism to identify WT or *irf8*^*−/−*^ animals.

For quantification of injury volume, the region of the PVZ that displayed signs of damage was manually outlined in ImageJ plane after plane. The volume was reconstituted using a custom script (Suppl. Files) and quantified using the plugin “3D geometrical measure” [48].

The Repair Index (RI) was estimated by calculating the following ratio: RI = 1-V24hpi/V4hpi.

### Trajectory analysis

MSD calculations were made with DiPer [49] using data obtained from MTrackJ. Trajectory characteristics such as straightness or z-displacement were manually extracted from MTrackJ data.

The Discrete Fréchet distance used to compare microglia and neuron trajectories in Fig. 3J was calculated with a custom script based on the Python library “trajectory distance”.

The trajectories anisotropy graph shown in Fig. 1I and was created using a custom script (Fig. S2, Suppl. Files). Displacement vector intersections from Fig. 3 were determined using a custom script (cf Suppl. Files).

### Image processing and analysis

The wound closure kinetics was estimated by measuring the fluorescence intensity in a region of interest delimiting the initial wound area in the PVZ over time. Long-term imaging data were registered for 3D motion in Fiji using the plugin correct 3D drift [50].

Before tracking, time-lapse images of Tg(h2a:GFP) were processed to improve the identification of each nucleus in Fiji using LocalNormalization [51] and denoising using the PureDenoise [52] plugins. Manual tracking of cells was done using the Fiji plugin MTrackJ [53].

The quantification of microglia accumulation kinetic was done by measuring the mean fluorescence intensity in the neuropil over time using ImageJ. The rostro-caudal extend curve was estimated by manually drawing a polyline starting rostrally and passing at an equal distance between the PVZ external and internal boundaries. Microglia accumulation positions were estimated from manually defined cell centroids in ImageJ after normalisation of the tectum to consider variations in size and position depending on the animal (Fig. S3). Before estimation of the repair index in Tg(Xla.Tubb:DsRed);Tg(irf8+/-) fish, images were blinded using a custom script (Suppl. Files). Image deconvolution for Fig. 5 was performed using DeconvolutionLab2 [54] in Fiji and a 2D calculated point spread function (PSF) using the Born-Wolf algorithm thanks to the PSF generator Fiji plugin [54]. Microglia morphology and endosome analysis were performed by manually drawing cell contours and measuring endosome diameters using Fiji.

### Laser severance and quantification

Cell process ablation in tectum-injured larval zebrafish Tg(her4.3:GFP-F);Tg(mpeg1:mCherry) was completed using a confocal microscope (SP8 inverse, Leica). Zebrafish larvae were mounted in 0.5% low melting point agarose as previously described in a multiwell cell culture imaging chamber (Ibidi) with the top of their heads touching the coverslip surface at 1 dpi. The imaging was completed in a temperature-controlled incubation chamber (25° C), using a 25x/0.95 HC FLUOTAR L water immersion objective (Leica). Single photo imaging was completed using a white light laser tuned to 488 nm (10% laser power) and 594 nm (2% laser power), with the emission range of the built-in HyD detectors set to the emission range of GFP and mCherry (respectively). Ablation was completed using a 5×1 µm region of interest (ROI) with a 2-photon laser tuned to 800 nm. All images were acquired at 14x zoom, with 10 planes (1 µm spacing). The ROI was exposed to the laser for all 10-plane images (512×512 pixels), with the focal point of the ROI placed on the astrocytic contact with the extended microglia during the full stack acquisition (15 sec). This was followed by a time series recording (5 min, 15 sec interval between frames, 512×512 pixels) to assess the movement of astrocytes and/or microglia. High-speed recordings were acquired for one optical section using the same imaging parameters, with a 0.65 ms interval between frames. Contacts between astrocyte-microglia contacts were selected based on the extension of astrocytic processes and microglia membranes. Laser-induced migrating microglia were those which were less than 10 µm away (2x the ablation ROI width) from the injury site at the time of laser activation. Astrocyte-astrocyte contacts were selected within the optic tectum adjacent to the injury site, but not touching it (circa 20 µm into the tectum from the internal edge of the injury site). For 3D images, the microglia and astrocytic processes were segmented manually in Fiji, using maximum intensity projections for 3D time series. The area change between frames was calculated from these ROI sets for either the microglia or astrocytes. For 2D images, the microglia were manually contoured using the built-in segmentation editor in FIJI [55]. The centre of the ablation ROI was calculated by applying a 5×1 µm ROI over the ablation site after rotating the image until the ablation site was horizontal. The relative edge distance of the microglia ROI to the centre point of the ablation site was quantified by comparing the centre of the ablation site to the leading edge of the microglia.

### Grey value distribution measurements for HCR signal

To measure the grey value distribution of the HCR signal within the lesioned tectum (Fig. 7), a single representative optical section (1 µm) was selected from a recorded confocal z-stack, encompassing expression within the lesion site. Pictures were converted into 8-bit greyscale and levels were adjusted by histogram clipping. Grey value distribution was measured along a 140 µm line (width 6 µm) originating at the tectal lesion site and extending within the unlesioned tectum. HCR signal grey values were measured using the Plot Profile tool in Fiji/ImageJ.

### Numerical simulations and programming frameworks

Simulations were performed using the PhysiCell framework [56] on a MacBook Pro (2.6 GHz 6-core i7 CPU, 16 Gb RAM) after compiling the custom C++ source code using GCC 11.0.3 (cf Suppl. Files).

All Python custom scripts were written using Spyder 4.1.2 and ran with Python 3.7.6 using the Anaconda3 environment.

### Statistical analysis

Curve fitting was done using the Curve Fitting Toolbox in MATLAB. t-Tests were performed using Excel (Microsoft). 2 and 3-factor nested ANOVA tests were performed according to (50) using Excel. Before estimation of the repair index in Tg(Xla.Tubb:DsRed);Tg(irf8+/-) fish, images were blinded using a custom script (Suppl. Files).

## ACKNOWLEDGEMENTS

We thank David Greenald (CRH, University of Edinburgh) and Katy Reid (CDBS, University of Edinburgh) for the kind gift of transgenic fish. We thank Jason Early (CDBS, University of Edinburgh) for his help with fluorescence microscopy. We thank Themistoklis M. Tsarouchas (SCRM, University of Edinburgh) for his help with genotyping. We thank Paul McKlin and Randy Heiland (Indiana University) for their help with the use of PhysiCell. We thank the uCreate Maker Space from the University of Edinburgh for 3D printing. This work was supported by the CMCB Light Microscopy Facility, a core facility of the CMCB Technology Platform at the Technische Universität Dresden. The laser ablation experiments and HCR imaging were performed on a Leica SP8 and the Andor Dragonfly (respectively) of the CMCB Light Microscopy Facility, a core facility of the CMCB Technology Platform at the Technische Universität Dresden. This work has been supported by the Biotechnology and Biological Sciences Research Council (BB/S0001778/1) (TB, CGB, FE), Moray Endowment Fund (FE), Alexander-von-Humboldt Professorship Award (CGB)

## Author Contributions (CRediT)

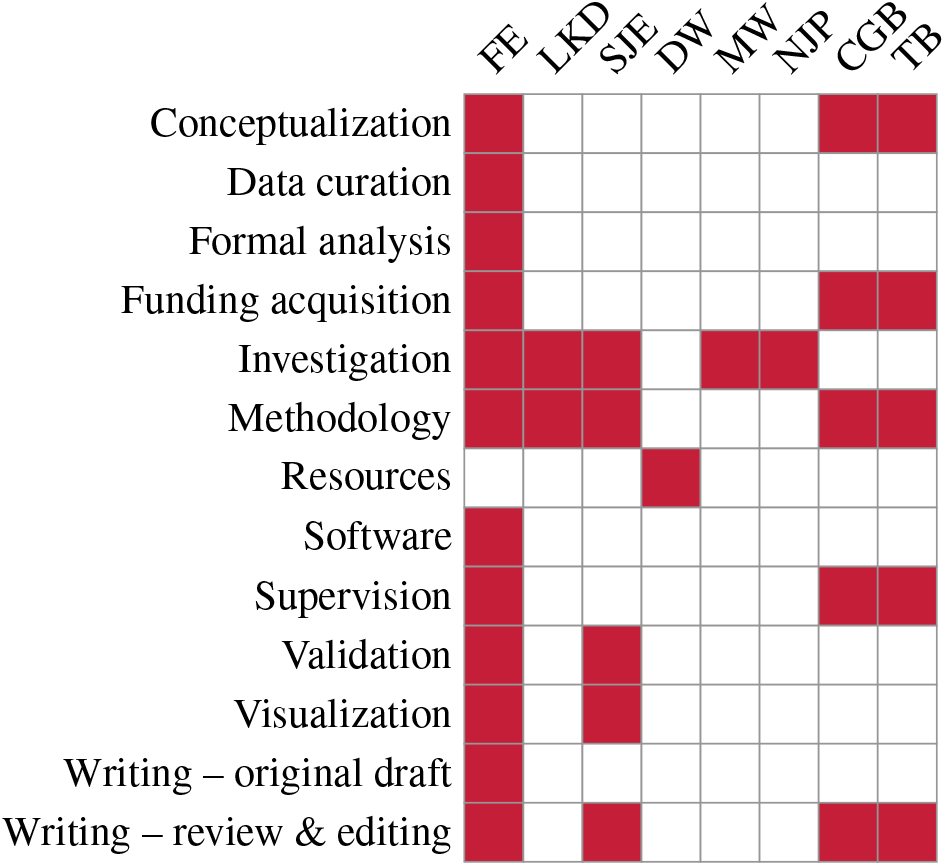

