## Supplemental Information for "Wound closure after brain injury relies on force generation by microglia in zebrafish"

#### Viscoelastic model of neurons dynamics (Fig. 2F, G).

We considered a model where neurons were attached to springs in a viscous medium. The position of a neuron can be obtained by solving Newton's first law of motion for a harmonic oscillator with an additional viscous force

$$m\,\frac{d^2x}{dt^2} + \eta\frac{dx}{dt} + k\,x = 0$$

Which gives:

$$x(t) = ae^{(-\nu t/2)}\cos\sqrt{(\frac{k}{m} - \frac{\nu^2}{4})}\,t - \varphi$$

We used this equation to fit the experimental fluorescence intensity curves shown in Fig. 1D using MATLAB.

#### Optic tectum multi-agent theoretical model.

The model we developed is based on the work from Ghaffarizadeh et al. [48]. We adapted the force balance and did not consider the interactions with the microenvironment. We also neglected cell cycle dynamics as it did not appear to play an essential role from our experimental observations (limited cell proliferation/neurogenesis).

The agents can exert/sense 4 forces:

1. Cell-cell adhesion
2. Cell-cell repulsion
3. Cell-cell elastic traction
4. Medium/ECM drag force

These forces are used in the computation in the form of gradients to update the velocity of each agent depending on its nature and position relative to every other agent. The skin agent has a particular role as it is not active or mobile. It was added to model the stiffness of the skin which affects cells and tissue dynamics and the overall forces balance. The shape of the PVZ was obtained from microscope image processing. Neurons were modelled as low mobility cells as observed experimentally on intact animals: a slow random motion without migration bias. Microglia were considered more motile, with an extra elastic force exerted between them and with neurons.

The elastic traction force is a phenomenological representation of forces exerted by microglia on astrocytic processes and the extracellular matrix. This allows for a simplified model able to reproduce the main observations with only a few parameters.

Neurons are represented only by their soma. We considered that the mechanical component of their protrusions is implicitly considered by the elastic traction force.

Below is the formulation of the agent velocity:

$$\vec{V_i} = \frac{1}{\eta}\sum_{j\in N(i)}\left[\vec{F}^{ij}_{cca} + \vec{F}^{ij}_{ccr} + \vec{F}^{ij}_{trac} + \vec{F}^{i}_{loc}\right]$$

Where $F_{cca}$ is the cell-cell adhesion force, $F_{ccr}$ the cell-cell repulsion force, $F_{loc}$, the locomotion force (for motile cells), and $F_{trac}$ the elastic traction force. $\eta$ is the medium effective viscosity.

The traction force is exerted only between microglia and other cells or between themselves. It follows Hooke's law and can be written as:

$$\vec{F}^{ij}_{trac} = \begin{cases} -k(\vec{r}_j - \vec{r}_i), & \text{if i,j is a microglia} \\ \vec{0}, & \text{if i,j is another types of cell ;} \end{cases}$$

#### Numerical implementation of the model.

We used the PhysiCell (version 1.7.1) framework for all simulations (http://PhysiCell.MathCancer.org). The program flow is detailed in [48].

Briefly, the velocity is updated for each agent, depending on its nature, its position, the forces exerted and the position of other agents interacting with it.

The table below summarizes the parameters used for the different agents in the simulations for the present work.

| Parameter | Neurons | Microglia | Skin cells | Biophysical meaning |
|---|---|---|---|---|
| Persistence time (min) | 10 | 10 | 0 | Directionality persistence of mobile cells |
| Migration speed (micron/min) | 0.01 | 1 | 0 | Cell velocity (related to Floc) |
| Relative repulsion | 5 | 5 | 5 | Cell-cell repulsion force amplitude |
| Relative adhesion | 0.1 | 0 | 0 | Cell-cell adhesion force amplitude |
| Elastic coefficient(1/min) | 5.10-7 | 5.10-7 | 5.10-7 | Amplitude of the elastic forces (Ftrac) |

**Table S1.** Parameters used for the simulations

The distribution of the different cell types was determined from fluorescence microscopy images using manual delimitation of the tectum, PVZ, and skin outline.

The positions of cells were generated using a Python script computing close-packing disc distribution inside an arbitrary 2D geometry.

For neurons, the actual shape of the PVZ was used. For skin cells, the thickness was artificially increased to mimic the skin stiffness. For microglia, positions were set manually by drawing ROI on the image mask in the neuropil region.

The generated files were used as inputs for the simulation program which generated a series of images at different time points. We used those images to generate the simulated wound closure kinetics in Figure 4C.

### Supplemental figures

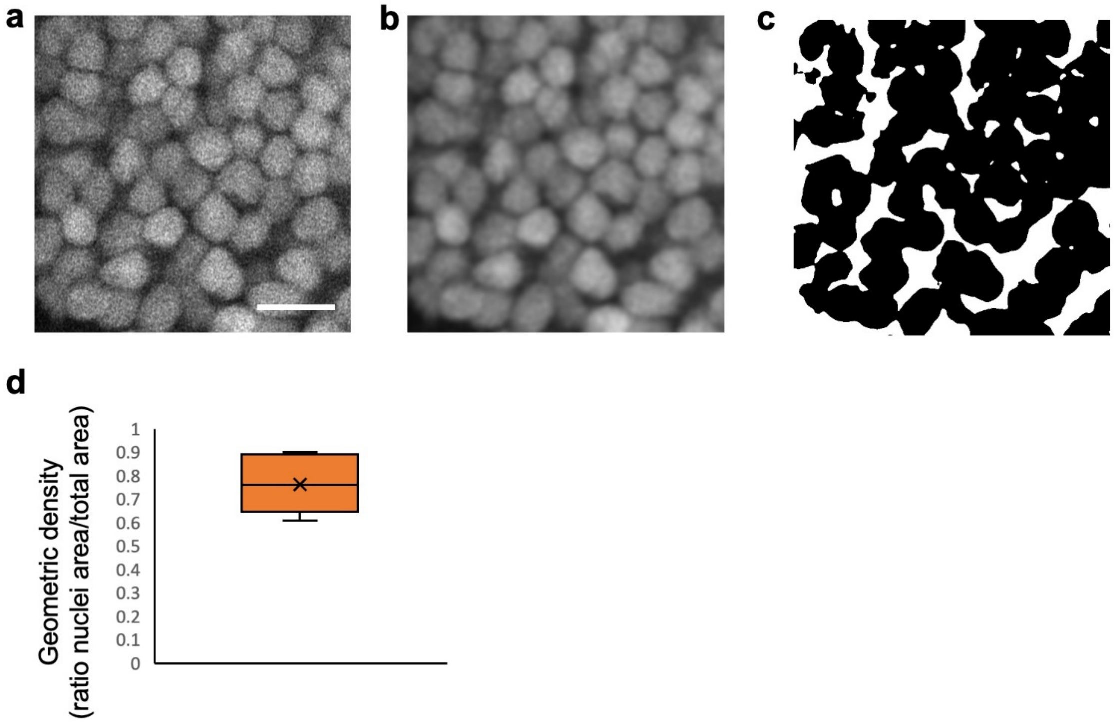


**Fig. S1. Neuronal nuclei show a dense hexagonal packing in the optic tectum** A Zoom-in on neuronal cell nuclei arrangement in the PVZ (Scale bar: 10 µm). B The same image after filtering using a Gaussian blur to reduce noise before quantification is shown. C. Thresholded binary images with nuclei in black and interstitial space in white are shown. D, Quantification of the geometrical density of neuronal cell nuclei (n = 6). Box plots show the median, box edges represent the 25th and 75th percentiles, and whiskers show the full data range.

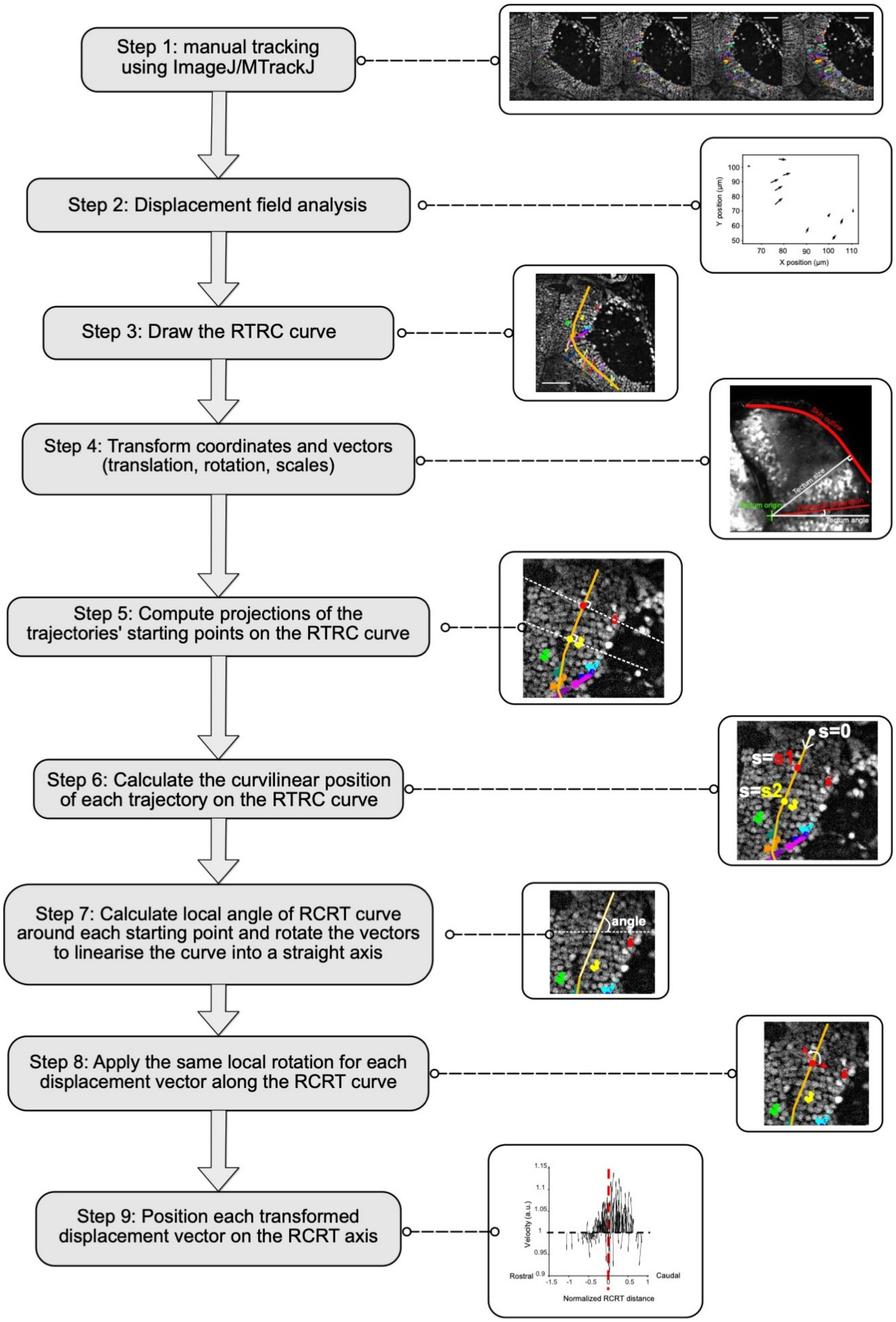


**Fig. S2. Workflow for the analysis of anisotropy of cell trajectories.** Analysis workflow used to generate the graphs Fig. 1I and Fig. 4G.

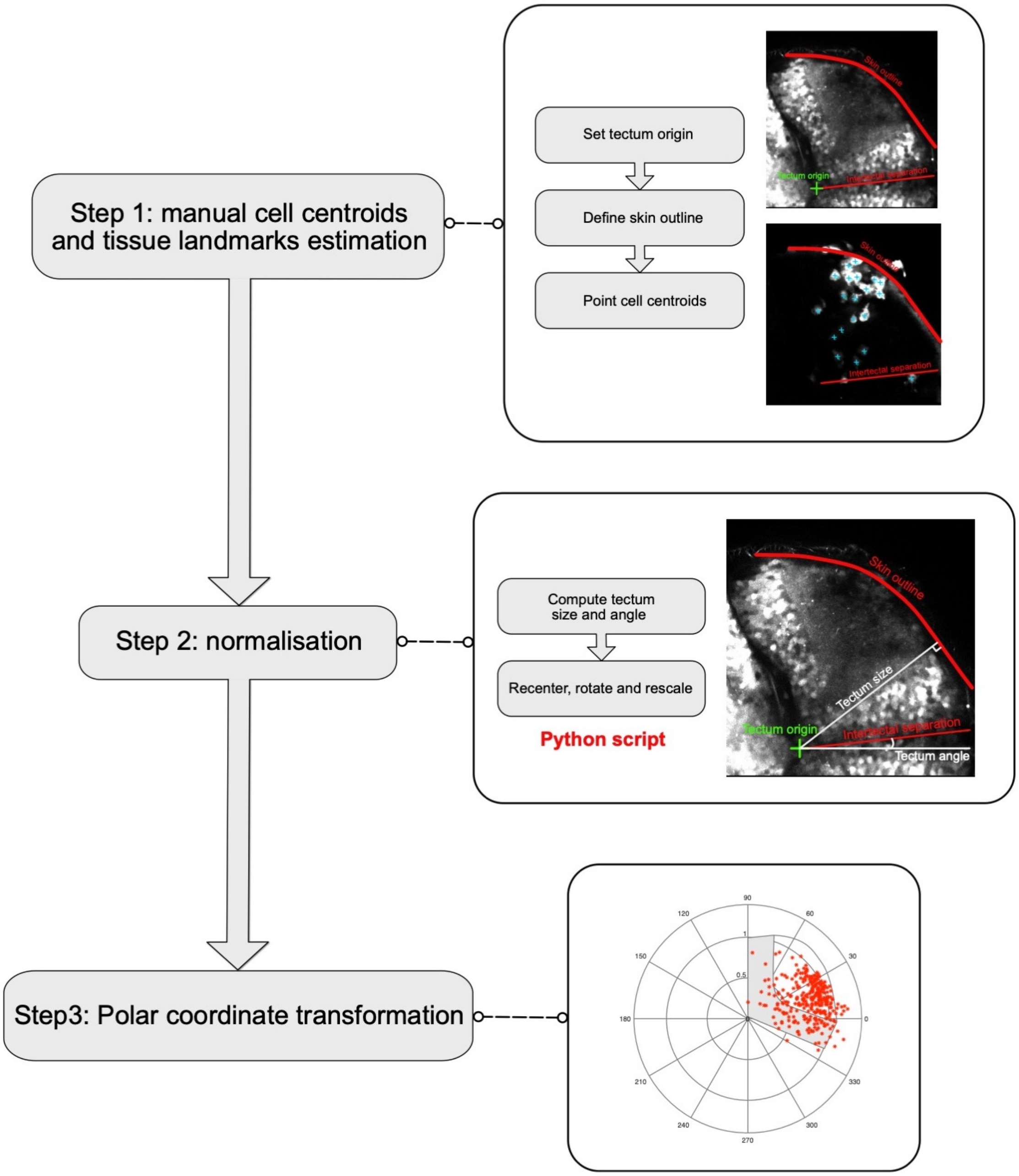


**Fig. S3. Microglia accumulation analysis workflow** Analysis workflow used to generate the graph in Fig. 3F.

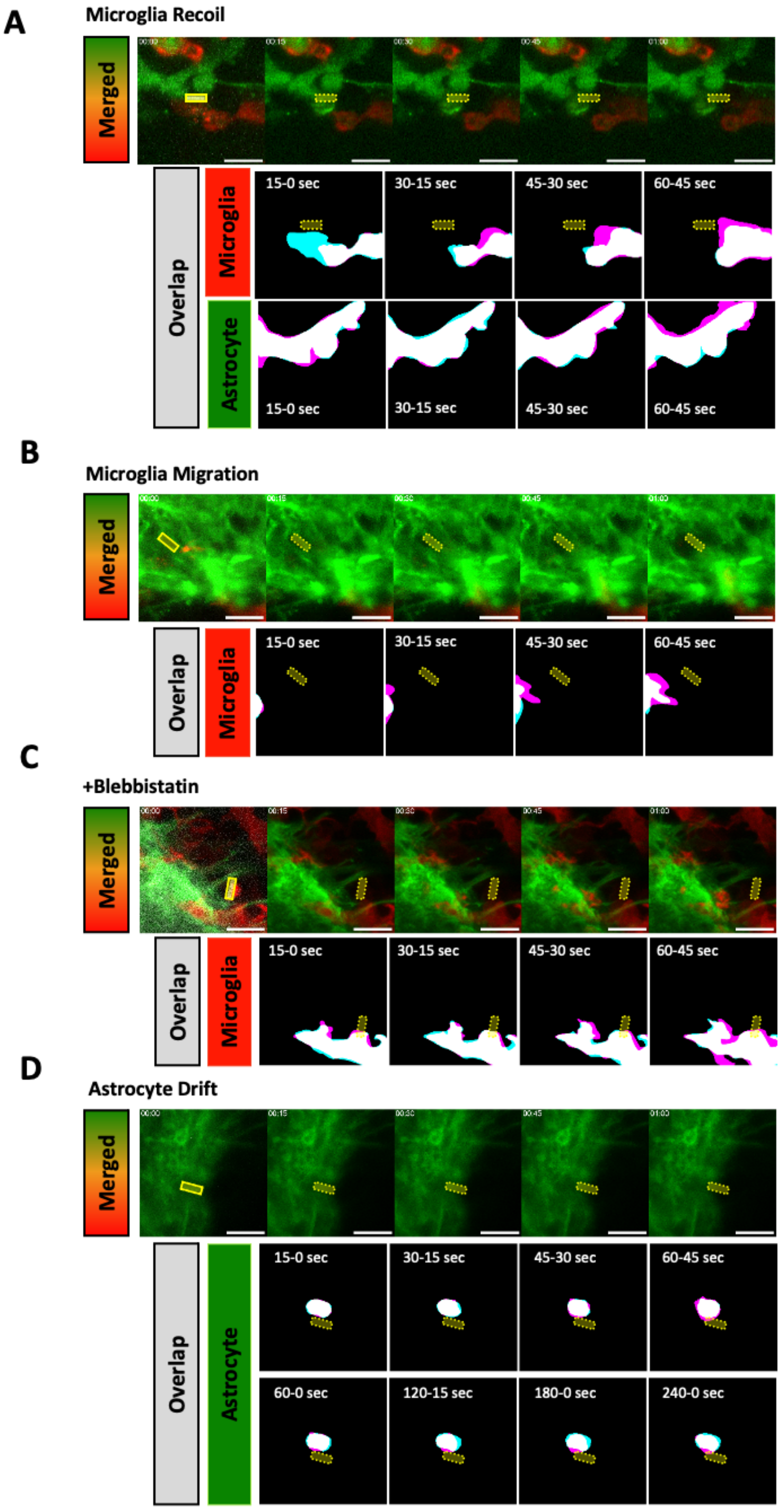


**Fig. S4. Additional examples and control conditions for cutting of microglia and astrocytic process contacts.** Dual channel (upper) and mask overlay (lower) montages for the first-minute post-cutting of recoiling microglia and the corresponding astrocytic process (A), a migrating microglia cell moving towards the cut site within the astrocytic processes (B), a contact between microglia and astrocytic process during blebbistatin treatment (C), and homotypic astrocytic process contacts (D). Overlays between adjacent time frames, first (cyan) and second (magenta, respectively). All scale bars represent 10 µm.

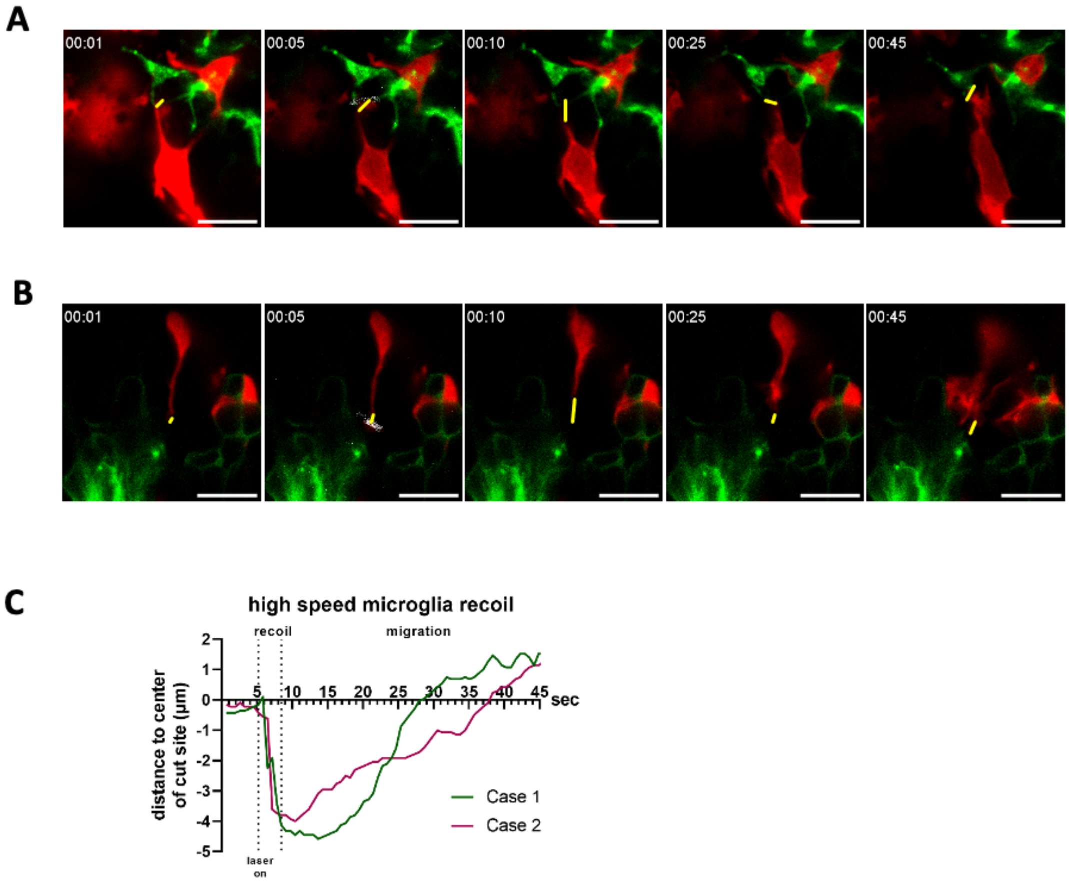


**Fig. S5. Single-plane high temporal resolution imaging of recoiling microglia contacts following laser cutting.** Individual montages of time-lapse images of two cases of microglia retraction in response to planar cutting of contacts with astrocytic processes (A, B). The distances between the centroid of the cut site and the edge of the microglia are labelled with yellow lines. Measurements of the distance between the leading edge of the microglia and the centroid of the laser cut site for each case over time (C). All scale bars represent 10 µm.

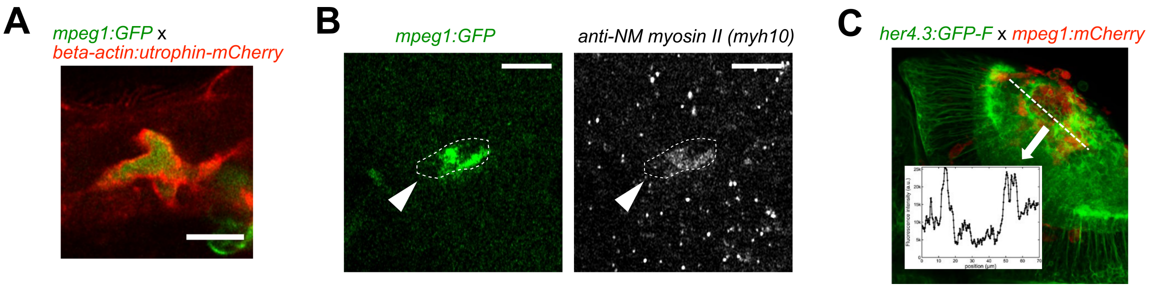


**Fig. S6. Microglia possesses the molecular machinery to exert forces and accumulation leads to increased GFAP detectability.** (A) A confocal micrograph of a microglia cell in the optic tectum which expresses both mpeg1:GFP and beta- actin:utrophin-mCherry, illustrating that the F-actin filaments accumulate along the exterior membrane of the microglia following stab injury. (B) A microglia cell (mpeg1:GFP) immunohistochemically labeled for non-muscle myosin II, the target for blebbistatin. The cell shows notable accumulation of myosin II. (C) A horizontal section of the tectum at 12 h post-injury is shown, together with a density plot for GFP fluorescence. Astrocytic processes labeled by her4.3:GFP-F show increased fluorescence next to the accumulation of microglia in the tectal neuropil. Scale bar in A & B = 10 µm.

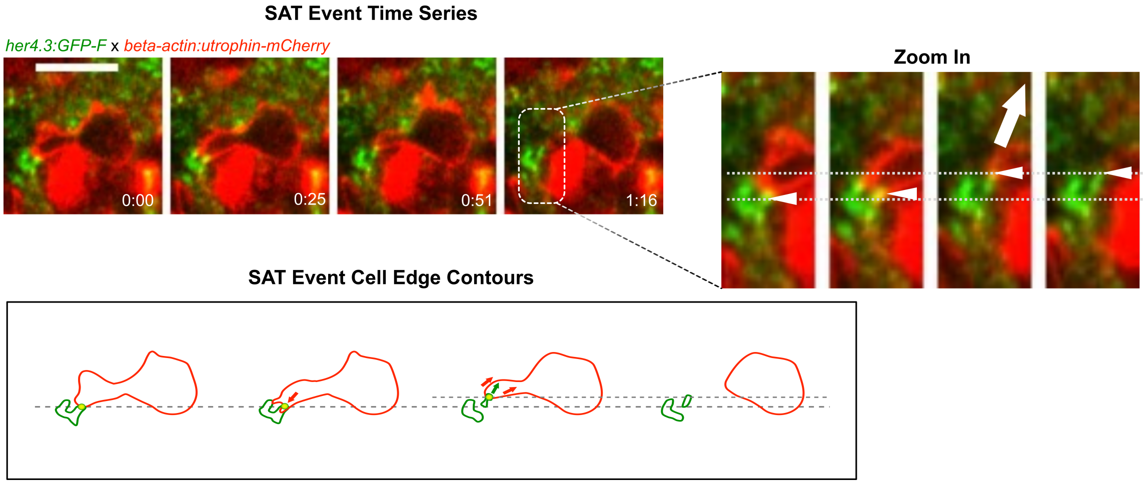


**Fig. S7. Utrophin labelling is enriched at sites of astrocyte pulling.** (Upper left) Fast time-lapse imaging sequence on a Tg(her4.3:GFP-F;beta-actin:utrophin-mCherry) larva showing a SAT process. The white arrow points out the adhesion event. (Right) Cropped sequence from upper left. White arrows pointing to astrocytic node pulled by microglia protrusion. The dashed line indicates the initial position. (Lower left) outline of microglia (red) and astrocytic structures (green) from images in the panel above. Arrows show the direction of displacement. Scale bar = 10 µm

Icp1

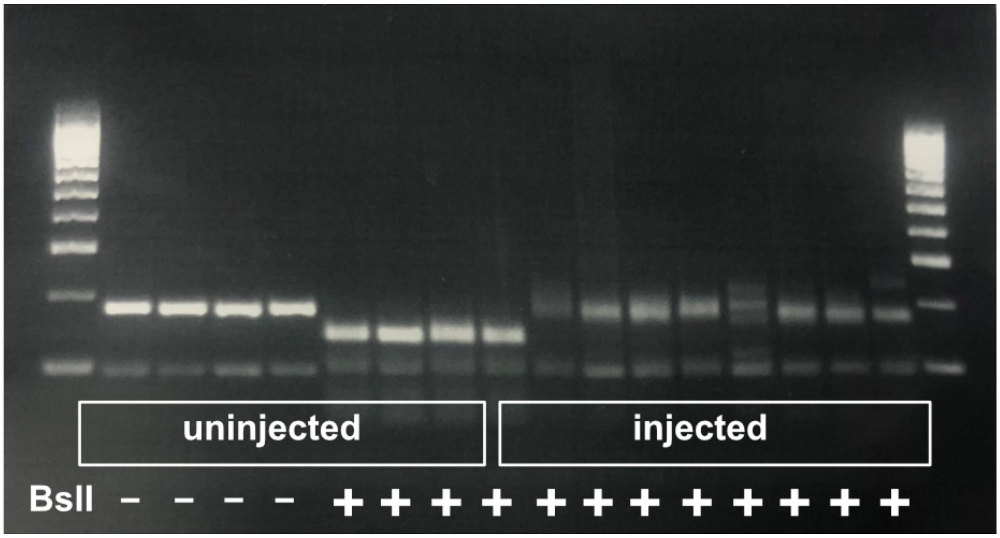


**Fig. S8. Determination of haCRs for Icp1.** Results of the RFLP technique on uninjected and injected animals for the targeted gene. Note that uninjected embryos show complete digestion with the indicated restriction enzymes, whereas haCR-injection efficiently alters the enzyme recognition site and prevents digestion.

### Supplemental movies

**Accessible through.** https://datashare.tu-dresden.de/s/XN6c7JJrAj75wSg

**Movie S1.** Time-lapse recording of a Tg(H2A:GFP) injured fish showing the kinetics of wound closure and the displacement of neuron cell bodies with individual trajectories (colored tracks). (Time interval between frames: 15 min. Scale bar: 20 µm)

**Movie S2.** Time-lapse recording of a Tg(XlaTubb:DsRed;mpeg1:GFP) injured fish showing the recruitment of microglia during the first 24h. (Time interval between frames: 30 min. Scale bar: 50 µm)

**Movie S3.** Example of numerical simulation of brain tissue repair using our multi-agent model. Blue: microglia, red: neurons, black: skin cells. (Time interval between simulated frames: 15 min.)

**Movie S4.** Maximum intensity projection movie of a Tg(her4.3GFP-F;mpeg1:mCherry) injured fish showing the increase in density of astrocytic processes around the accumulated microglia during the repair process. (Time interval between frames: 15 min. Scale bar: 50 µm)

**Movie S5.** Time-lapse recording of a Tg(her4.3GFP-F;mpeg1:mCherry) injured fish showing an example of microglia exerting traction on astrocytic processes. (Time interval between frames: 25 s.)

**Movie S6.** Time-lapse recording of a Tg(her4.3GFP-F;mpeg1:mCherry) injured fish showing an example of a migration event where astrocytic processes are pulled by a moving microglia. (Time interval between frames: 6 s.)

**Movie S7.** Time-lapse recording of a Tg(her4.3GFP-F;mpeg1:mCherry) injured fish showing an example of microglia "knitting" astrocytic processes. (Time interval between frames: 25 s. Scale bar: 10 µm)

**Movie S8.** Time-lapse recording of a Tg(Her4.3GFP-F; utrophin:mCherry) injured fish showing an example of microglia exerting traction on astrocytic processes correlated with a recruitment of the actin cytoskeleton as the traction point. (Time interval between frames: 25 s. Scale bar: 10 µm)

**Movie S9.** Time-lapse recording of a Tg(her4.3GFP-F;mpeg1:mCherry) injured fish showing an example of microglia recoil in response to laser severance of the cell's contact with an astrocytic process. The video is a maximum intensity projection of a 3D image (10 µm total depth, 1 µm plane intervals. Time interval between frames: 15s. Scale Bar: 10 µm)

**Movie S10.** Time-lapse recording of a Tg(her4.3GFP-F;mpeg1:mCherry) injured fish showing an example of microglia recoil in response to laser severance of the cell's contact with an astrocytic process. The video is a 2D image of a single optical plane (Time interval between frames: 0.65s. Scale Bar: 10 µm)
